# The replication properties of a contemporary Zika virus from West Africa depends on NS1/NS4B proteins

**DOI:** 10.1101/2024.03.14.584947

**Authors:** Dana Machmouchi, Marie-Pierre Courageot, Chaker El-Kalamouni, Alain Kohl, Philippe Desprès

**Affiliations:** Processus Infectieux en Milieu Insulaire Tropical (PIMIT), Université de La Réunion, INSERM U1187, CNRS 9192, IRD 249, Plateforme Technologique CYROI, 97490 Sainte-Clotilde, La Réunion, France; UR7506-BioSpect, Université de Reims Champagne-Ardennes, 51100 Reims, France; Liverpool School of Tropical Medicine, Pembroke PI, Liverpool L3 5QA, UK

**Keywords:** Arbovirus, Zika virus, genetic reverse, viral replication, nonstructural protein, mutant virus, chimeric virus, host-cell interactions, antiviral innate immunity, stress granules

## Abstract

Zika virus (ZIKV) have become a global health problem over the past decade due to the extension of the geographic distribution of ZIKV of Asian genotype. Epidemics of Asian ZIKV have been associated with developmental disorders in humans. ZIKV of African lineage would have an epidemic potential associated to fetal pathogenicity requiring a greater attention towards the most recently isolated viral strains from West Africa. In the present study, an infectious molecular clone GUINEA-18 has been obtained from viral strain ZIKV-15555 that had been sequenced from an individual infected by ZIKV in Guinea in 2018. A molecular clone-based comparative study between GUINEA-18 and viral clone MR766^MC^ from historical African ZIKV strain MR766 revealed a lower replication rate for GUINEA-18 associated to a weaker cytotoxicity and reduced innate immune system activation in Vero E6, A549 and HCM3 cell lines. Analysis of chimeric viruses between MR766^MC^ and GUINEA-18 stressed the importance NS1/NS4B proteins with a particular focus for NS4B on GUINEA-18 replication properties. ZIKV has developed strategies to prevent cytoplasmic stress granule formation which occurs in response to virus infection. Study of G3BP protein showed that GUINEA-18 but not MR766^MC^ was efficient to inhibit stress granule assembly in A549 cells subjected to a physiological stressor. GUINEA-18 depends on NS1/NS4B proteins for suppressing stress granule response to environmental stress. The involvement of GUINEA-18 NS1/NS4B proteins on virus replication capability and host-cell responses to ZIKV infection raises the question of the importance of nonstructural proteins in the pathogenicity of contemporary viral strains from West Africa.

**AUTHOR SUMMARY:** Most of studies having for objectives to understand the biology of Zika virus (ZIKV) were carried out using epidemic viral strains of Asian lineage. It is now admitted that ZIKV of African genotype would have also a great epidemic potential associated a high risk of fetal pathogenicity. Today, it is urgent to improve our knowledge on recently isolated ZIKV strains in West Africa. In our study, we used the sequence of viral strain from an individual infected by ZIKV in Guinea in 2018 to generate an infectious molecular clone. Analysis of viral clone highlighted the preponderant role of NS1/NS4B proteins in virus replication strategy and cell interactions with a particular focus on ZIKV-specific stress granule formation blockade. We believe that our data will improve our knowledge on the biology of contemporary West Africa ZIKV opening perspectives towards a better understanding on the pathogenicity of African viral strains.

## INTRODUCTION

Zika virus (ZIKV) infection is widespread medically important mosquito-borne viral disease for which there are not vaccines nor curative treatments (1). ZIKV circulates in Africa and Asia and more recently in Americas, where an enzootic transmission cycle appears to involve primates, with the mosquitoes *Aedes aegypti* and less competent *Ae*des *albopictus* as potential vectors (1,2). ZIKV strains are phylogenetically classified as African, Asian and now Asian/American genotypes (3). In the past decade, there has been unexpected expansion of the geographic distribution of ZIKV strains of Asian lineage and their rapid spread caused major epidemics in the South Pacific in 2013 and for the first time in Americas in 2015. Emergence of Asian ZIKV strains has been associated with a polyradiculoneuropathy (Guillain-Barré syndrome) and unprecedented severe complications in humans grouped together under the umbrella term Congenital Zika Syndrome (CZS) with teratogenic effects presenting as microcephaly and other serious neurological abnormalities (1,3,4,5). Although ZIKV transmission in humans classically involves a blood meal by infected female mosquitoes, sexual contact, blood transfusion and intrauterine transmission have been documented as non-vectored transmission routes (1). ZIKV is an enveloped, positive, single-stranded RNA virus belonging to orthoflavivirus genus of *Flaviviridae* family (1,2). The genomic RNA of about 11,000 nucleotides of length contains a single open reading frame flanked by highly structured 5’ and 3’ non-coding regions (NCRs) (1,2). The 5’NCR which is essential for initiation of viral RNA translation, includes two stem-loop (SLs) which are required as promoter of viral replication and for genome cyclization. The 3’NCR is organized in several domains with a variable structural element followed by number of conserved SLs just downstream from the translation stop codon. The genomic RNA is translated in a large polyprotein precursor that is co- and post-translationally processed into three structural proteins, capsid (C), precursor membrane (prM/M) and envelope (E) protein followed by seven nonstructural protein NS1, NS2A, NS2B, NS3, NS4A, NS4B and NS5 (1,2). Structural proteins C, prM, and E are required for the formation of infectious viral particles whereas NS proteins play important roles in viral RNA replication, protein processing, and virion assembly (1,2). The NS proteins also contribute to innate immune subversion strategies of ZIKV (6). NS1 exists as intracellular homodimer and secreted soluble hexamer which is associated with different types of lipid molecules (7). Membrane-associated NS2A is involved in RNA replication and virion assembly (8–10). NS3 protein is an enzyme with both serine protease and NTPase/helicase activities (11). The ER resident NS4A protein is an essential component of viral replication complexes (VRCs) playing a role in virus-induced membrane reorganizations (12). Membrane bound NS4B protein contributes to VRCs and host-immunomodulation (13). Lastly, the large NS5 protein consists of an N-terminal MTase and a C-terminal RNA-dependent-RNA polymerase (14).

Recently isolated West Africa ZIKV strains have been recently identified as potentially highly teratogenic in humans demanding urgent attention in public health (15,16). Also, ZIKV strains of African lineage were found to display high levels of transmission by *Ae.albopictus* pointing at their elevated risk of introduction in regions where the mosquito vector has established increasing populations (17,18). Such recently uncovered features of ZIKV of African lineage are worrying and require further investigations. The risk assessment associated to African ZIKV lineage virus implies to improve our knowledge on the virological characteristics of recently isolated viral strains circulating in sub-Saharan Africa. In 2018, ZIKV-15555 genomic RNA sequence (Genbank accession number MN025403) has been obtained from an individual infected by ZIKV in Guinea. Viral strain ZIKV-15555 shares a high degree of sequence homology with historical African ZIKV strain MR766 (Genbank accession number LC002520) isolated from a non-human primate in Uganda in 1947. MR766 has essentially retained its genetic integrity even though the virus has been initially considered as a widely laboratory-adapted viral strain. This supports the idea that African ZIKV lineage viruses would be genetically more stable than it had been previously anticipated. An infectious molecular clone of MR766-NIID (hereafter named MR766^MC^) was obtained by reverse genetic using the ISA method (19,20). In an effort to better understand the virological features of contemporary ZIKV strains of African lineage, we have generated an infectious molecular clone of ZIKV-15555 referred hereafter as GUINEA-18. A comparative analysis between GUINEA-18 and MR766^MC^ in different cell lines have shown differences in viral replication rate and host-cell responses to virus infection. Our data indicated that replication strategy of GUINEA-18 depends on NS1/NS4B proteins.

## MATERIALS and METHODS

### Cells and antibodies

Human embryonic kidney HEK-293T (ATCC, CRL-1573), human carcinoma epithelial lung A549 (Invivogen Inc, Toulouse, France), human microglial clone 3 HCM3 (ATCC, CCL-3304 from Dr N. Jouvenet, Institut Pasteur, Paris, France) and monkey kidney normal Vero E6 (CCL-81, ATCC, Manassas, VA, USA) cells were grown in Dulbecco’s modified Eagle’s medium (DMEM) growth medium supplemented with 10 or 5% of heat-inactivated fetal calf serum (FBS, Dutscher, Brumath, France), respectively, and antibiotics (PAN Biotech Dutscher, Brumath, France) at 37 °C under a 5% CO_2_ atmosphere. The mouse anti-*pan* orthoflavivirus envelope E protein monoclonal antibody (mAb) 4G2 was produced by RD Biotech (Besançon, France). Donkey IgG anti-mouse IgG Alexa Fluor 488 or Alexa Fluor 594 and donkey IgG anti-rabbit IgG Alexa Fluor 594 secondary antibodies were purchased from Invitrogen (Thermo Fisher Scientific, Illkirch-Graffenstaden, France). Goat IgG anti-mouse IgG-horseradish peroxidase (HRP) conjugated were purchased from Abcam (Cambridge, UK). Blue-fluorescent DNA stain 4’,6-diamidino-2-phenylindole (DAPI) was purchased from Euromedex (Souffelweyersheim, France). Lipofectamine 3000 (Thermo Fisher Scientific, Illkirch-Graffenstaden, France) was used for transfection, according to the manufacturer’s instructions.

### Design of infectious ZIKV molecular clones and chimeric viruses

The infectious molecular clone MR766^MC^ derived from historical African 1947 ZIKV strain MR766-NIID (Genbank accession number LC002520) was previously described by Gadea *et al.* (2016) (20). The infectious molecular clone GUINEA-18 derived from viral strain ZIKV-15555 (Genbank accession number MN025403) was produced as described for MR766^MC^. Because the sequences of non-coding regions (NCRs) deposited in Genbank were lacking for the first nucleotides of 5’NCR and the last nucleotides of 3’NCR, the extremities of MR766^MC^ sequence were chosen to complete the 5’ and 3’ NCRs of GUINEA-18. Based on the genetic reverse method ISA (19), the design of GUINEA-18 genomic RNA into three fragments Z-1^GUINEA-18^, Z-23^GUINEA-18^ and Z-4^GUINEA-18^ was chosen to mimic Z-1^MR766^, Z-23^MR766^ and Z-4^MR766^ used to construct MR766^MC^.The fragment Z-1^GUINEA-18^ includes the CMV promoter immediately adjacent to the 5’NCR followed by the coding region for residues 1 to 712 of viral polyprotein (C, prM and 80% E). The Z-23^GUINEA-18^ fragment encompasses the residues 702 to 2684 (20% E followed by NS1 to NS4B) of viral polyprotein. The fragment Z-4^GUNEA-18^ gene corresponds residues 2674 to 3423 of viral polyprotein (mostly NS5) followed by the 3’NCR followed by a hepatitis delta virus ribozyme and ended by a SV40 poly(A) signal. The synthetic Z-1^GUINEA-18^, Z-23^GUINEA-18^ and Z-4^GUINEA-18^ genes were synthetized and inserted into plasmid pUC57 by Genecust (Boynes, France). The Z-1^GUINEA-18^, Z-23^GUINEA-18^ and Z-4^GUINEA-18^ fragments were amplified by PCR from their respective plasmids using a set of specific primers (Table S1) so that Z-1^GUINEA-18^ and Z-23 ^GUINEA-18^ ^or^ Z-23^GUINEA-18^ and Z-4^GUINEA-18^ matched on at least 30 nucleotides. Also, Z-1^MR766^ and Z-23 ^GUINEA-18^ fragments as well as Z-23 ^GUINEA-18^ and Z-4^MR766^ fragments can match between them in order to generate chimeric viruses between MR766^MC^ and GUINEA-18. Site-directed mutagenesis on Z-23^MR766^ and Z4^MR766^ genes was performed by Genecust and the sequences verified by Sanger method.

### Recovering of infectious ZIKV molecular clones

The purified PCR products corresponding to Z-1^GUINEA-18^, Z-23^GUINEA-18^ and Z-4^GUINEA-18^ fragments were transfected in HEK-293T cells using Lipofectamine 3,000 (ThermoFisher, France). After 4 days, cell supernatants were recovered and used to infected Vero E6 cells in a first round of amplification (P1). After 3 to 5 days, P1 was recovered and amplified for another 2 to 3 days to produce a working virus stock P2. The resulting P2 virus was designed hereafter as GUINEA-2018. To produce chimeric ZIKV containing the structural protein region of MR766^MC^ followed by the nonstructural region of GUINEA-18, HEK-293T cells were transfected with PCR products containing Z1^MR766^, Z-23^GUINEA-18^ and Z-4^GUINEA-18^ sequences. To generate chimeric MR766^MC^ in which all or part of NS proteins were replaced by their counterpart of GUINEA-18, PCR was performed on Z-23^GUINEA-18^ fragment using specific primer pairs that were designed so that GUINEA-18 NS genes were inserted into MR766^MC^ backbone after transfection of HEK-293T cells. Virus stocks P2 grown on Vero E6 cells were used in this study. Virus stock titers in plaque forming unit per ml (PFU.mL^-1^) were determined by a standard plaque-forming assay as previously described (21).

### RT-qPCR

Total RNA was extracted from cells using RNeasy kit (Quiagen) and reverse transcription was performed using random hexamer primers (intracellular viral RNA) and MMLV reverse transcriptase (Life Technologies). Quantitative PCR was performed on a ABI7500 Real-Time PCR System (Applied Biosystems, Life Technologies, Villebon-sur-Yvette, France). Data were normalized using 36B4 as RPLP0 housekeeping gene. For each single-well amplification reaction, a threshold cycle (Ct) was calculated using the ABI7500 program (Applied Biosystems, Life Technologies) in the exponential phase of amplification. Relative change in gene expression was determined using the 2∂∂*C*t method and reported relative to the control. The primers used in this study are listed in Table S1.

### Recombinant NS4B protein

Mammalian codon-optimized genes coding for ZIKV 2K-NS4B (residues 2246 to 2527 from viral polyprotein) from viral strains MR766, ZIKV-15555, or epidemic Brazilian 2015 strain BeH819015 (Genbank accession number KU365778) were established using *Homo sapiens* codon usage as reference. A glycine-serine spacer followed by a FLAG tag were inserted in-frame at the C-terminus of recombinant NS4B protein. The synthesis of gene sequences and their cloning into *Nhe*-I and *Not*-I restriction sites of the pcDNA3.1-hygro (+) vector plasmid to generate recombinant plasmids pcDNA3/MR766.NS4B, pcDNA3/ZIKV-15555.NS4B, and pcDNA3/BeH819015.NS4B were performed by Genecust (Boynes, France). A deletion mutant coding for ZIKV-15555 NS4B protein without the 2K peptide (residues 2270 to 2527 from viral polyprotein) was cloned into pcDNA3.1-hygro (+) to generate pcDNA3/(Δ2K)-ZIKV-15555.NS4B. The plasmid sequences were verified by Sanger method. Endotoxin-free plasmid DNA purification was performed by Genecust (Boynes, France). A549 and HCM3 cells were transfected with plasmids using Lipofectamine 3,000.

### The eGFP-G3BP fusion protein

Mammalian codon-optimized gene coding for enhanced Green Fluorescent Protein (eGFP) carrying the A206K monomeric mutation (Genbank accession number AAB02572) followed by G3BP1 protein (Genbank accession number U32519) was established using *Homo sapiens* codon usage as reference. A glycine-serine spacer was inserted in-frame between eGFP and G3BP. The synthesis of gene sequence and cloning into *EcoR*-I and *Not*-I restriction sites of the pcDNA3.1-vector plasmid to generate recombinant plasmid pcDNA3/eGFP-G3BP was performed by Genecust (Boynes, France). The plasmid sequence was verified by Sanger method. Endotoxin-free plasmid DNA purification was performed by Genecust (Boynes, France). A549 cells were transfected with pcDNA3/eGFP-G3BP using Lipofectamine 3,000.

### Confocal immunofluorescence assay

Cells seeded on coverslips were fixed with 3.7% paraformaldehyde (PFA) in PBS at RT for 10 min. For analysis of eGFP-G3BP expression, cells were directly visualized by confocal fluorescence microscopy. For analysis of cells transfected with pcDNA3/eGFP-G3BP and then infected with ZIKV, fixed cells were permeabilized with nonionic detergent Triton X-100 at the final concentration of 0.1% in PBS. Cells were stained with anti-E mAb 4G2 in PBS containing 1% bovine serum albumin (BSA). Goat anti-mouse IgG Alexa Fluor 594 was used as secondary antibody. The capture of the fluorescent signal was allowed with a confocal fluorescence Nikon Eclipse TI2-S-HU microscope equipped with x63 objectives coupled to the imaging software NIS-Element AR (Nikon, Champigny-sur-Marne, France). Image processing based on FIJI/ImageJ software was performed to estimate the size in µm^2^ of eGFP-positive condensates.

### Flow cytometry assay

Cells were harvested after trypsinization and fixed with 3.7% PFA in PBS at RT for 10 min. A solution of Triton X-100 (0.15%) in PBS was used to permeabilize the fixed cells for 5 min. After incubation of cells with a blocking solution for 10 min, cells were stained with mouse anti-E protein mAb 4G2 or rabbit anti-FLAG antibody as primary antibody for 1h. Donkey IgG anti-mouse or anti-rabbit IgG Alexa Fluor 488 served as secondary antibody. Immunostained cells were subjected to a flow cytometric analysis using FACScan flow cytometer (CytoFLEX, Beckman Coulter, Brea, CA, USA). For each assay, at least 10,000 cells were analyzed and the percentage of positive cells was determined using CytExpert software (version 2.1.0.92, Beckman Coulter, Brea, CA, USA).

### Immunoblot assay

Cells were lysed with RIPA lysis buffer and clarified cell lysates were separated by in-house 12% SDS–PAGE gel. Proteins were transferred onto nitrocellulose NC Protran membrane and after blocking with 5% no-fat dry milk in PBS-Tween for 30 min, the membrane was probed with rabbit anti-FLAG antibody for 1h at room temperature (RT). Anti-rabbit IgG-horseradish peroxidase (HRP) conjugate was used as secondary antibody. Membranes were incubated with Pierce ECL Western blotting substrate (Thermo Fisher Scientific) and exposed using Amersham imager 680 (GE Healthcare).

### Cytotoxic assay

Cell damages were evaluated measuring lactate dehydrogenase (LDH) release. Supernatants of infected cells were recovered and subjected to a cytotoxicity assay, performed using CytoTox 96® non-radioactive cytotoxicity assay (Promega, Madison, WI, USA) according to manufacturer instructions. Absorbance of converted dye was measured at 490 nm (Tecan). Results of LDH activity are presented with subtraction of control values.

### Statistical analysis

All statistical tests were done using the software GraphPad Prism version 10.1.1. Unpaired *t* test and ANOVA were used in this study.

## RESULTS

### Characterization of infectious molecular clone GUINEA-18

Contemporary West Africa ZIKV strain ZIKV-15555 that had been sequenced from an individual infected by ZIKV in Guinea in 2018 differs from historical African ZIKV strain MR766 by 49 amino acid substitutions into polyprotein (Table 1). There also 2 and 8 mutations between the 5’ and 3’ NCRs of ZIKV-15555 and MR766, respectively.

**Table 1.**
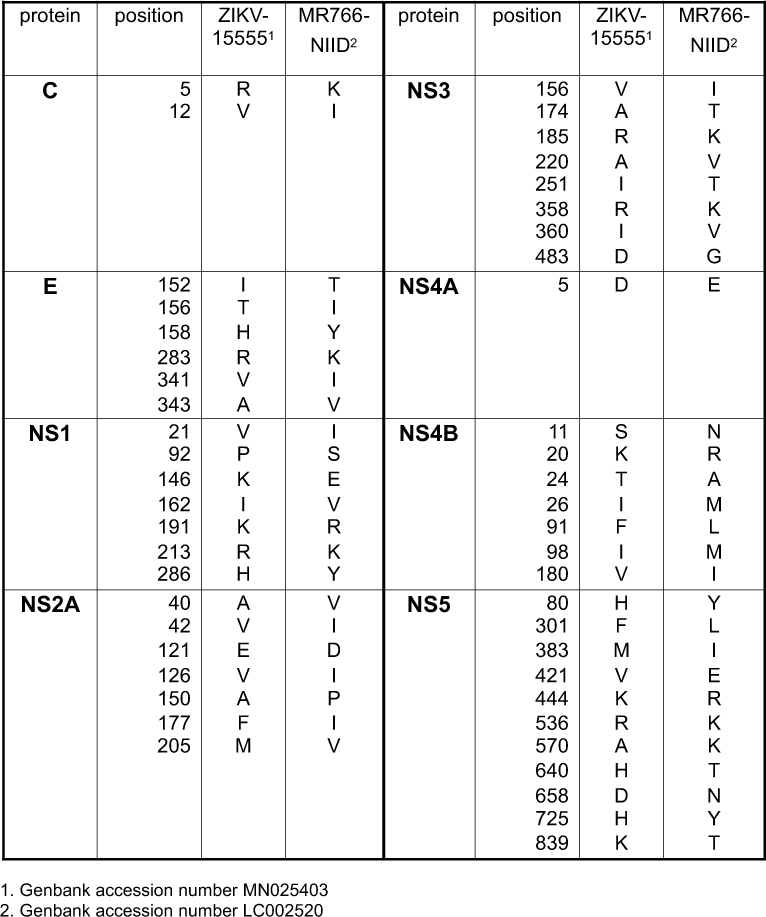
Amino acid changes between African ZIKV strains ZIKV-15555 (Genbank access number MN025403) and MR766-NIID (Genbank accession number LC002520).

Our previous works on African ZIKV involved the infectious clone MR766^MC^ which was achieved by constructing a molecular clone of MR766-NIID using the ISA method (19). Similarly, a molecular clone GUINEA-18 has been obtained by constructing an infectious clone derived from ZIKV-15555 genomic RNA (Fig. 1A). The complete genome of ZIKV-15555 was assembled in transfected HEK-293T cells virus progeny was twice amplified on Vero E6 cells. For this study, we used ZIKV-15555 sequences which overlap with their counterparts from MR766^MC^ in order to generate chimeric viruses between the two infectious molecular clones (Fig. 1A). We noted that GUINEA-18 and MR766^MC^ generated plaques of medium size on Vero E6 cells (Fig.1A).

**Figure 1.**
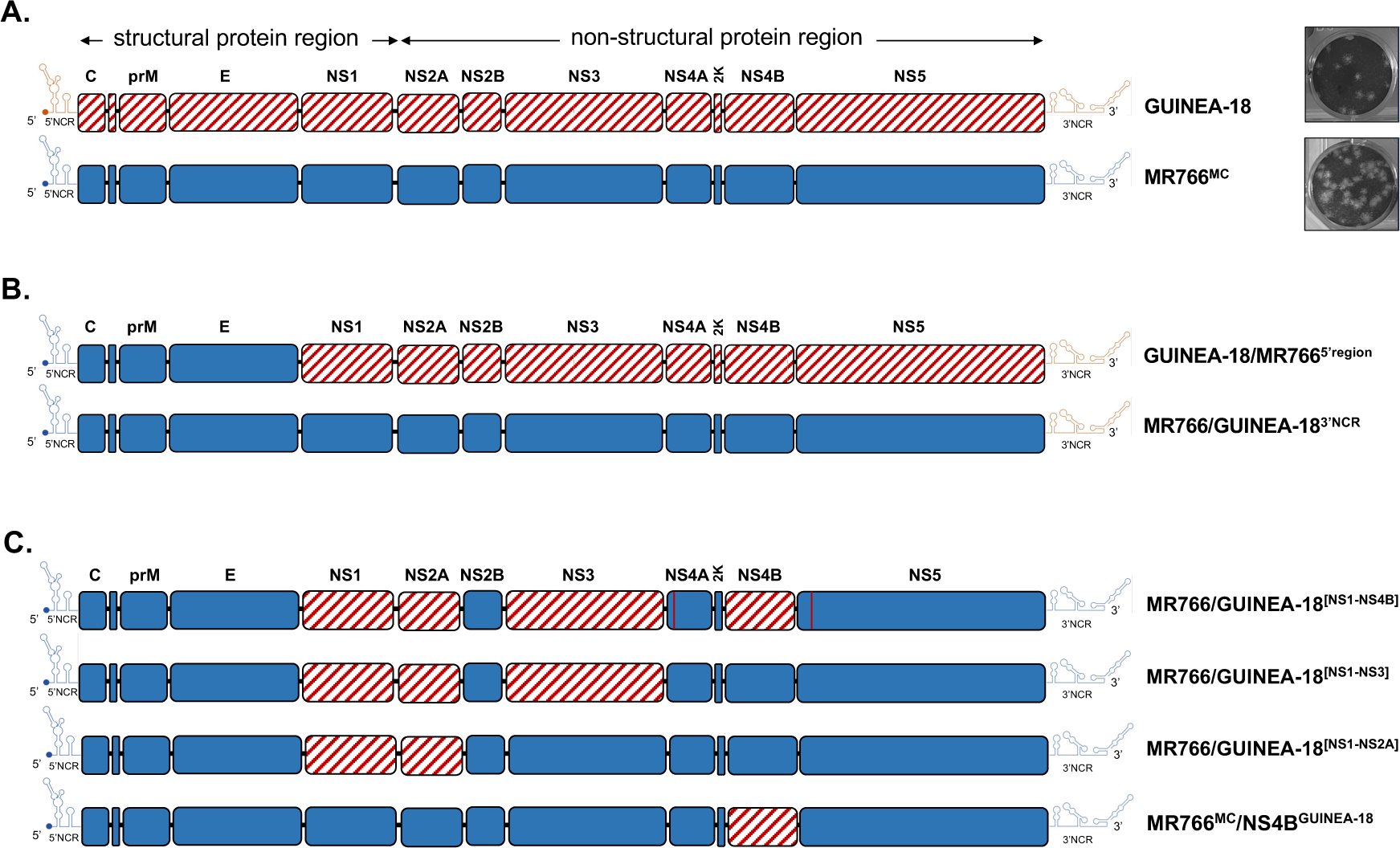
ZIKV molecular clones GUINEA-18, MR766 and chimeric viruses. Generation of chimeric ZIKV molecular clones are shown along with their parental clones. The chimeric viruses were made between the MR766 (blue) and GUINEA-18 (red hatched) ZIKV molecular clones. *In* (**A**), schematic representation of viral clone GUINEA-18 derived from ZIKV strain ZIKV-15555. The production of MR766^MC^ has been described by Gadea *et al*., (2016). The ZIKV structural and nonstructural proteins are indicated. NCR: non-coding region. In right, virus plaques on Vero E6 cells are shown. *In* (**B**), GUINEA-18/MR766^5’region^ is a chimeric virus containing the 5’NCR and structural protein region of MR766^MC^ into GUINEA-18 backbone. MR766/GUINEA-18^3’NCR^ is a chimeric virus containing the 3’NCR of GUINEA-18 into MR766^MC^ backbone. *In* (**C**), MR766/GUINEA-18^[NS1-NS2A]^, MR766/GUINEA-18^[NS1-NS3]^, MR766/GUINEA-18^[NS1-NS4B]^ and MR766/GUINEA-18^[NS4B]^ are chimeric viruses with different GUINEA-18 NS genes inserted into MR766^MC^ backbone. conserved NS2B protein between MR766 and GUINEA-18 is colored in blue in chimeric viruses. The amino-acid substitutions NS4A-E5D and NS5-N11S that differentiate MR766^MC^ from GUINEA-18 (Table 1) are indicated by red vertical lines.

The GUINEA-18 and MR766^MC^ E proteins differ by the amino-acid substitutions I152T/T156I/H158Y that are part of ZIKV glycan loop (GL) region. The mutation E-T156I in MR766^MC^ causes a loss of the carbohydrate attachment site on residue N154 leading to non-glycosylated protein. Immunoblot assay using anti-E mAb 4G2 showed that GUINEA-18 E protein migrated slower than MR766^MC^ E protein (Fig. S1). The migration profile of GUINEA-18 E protein is consistent with a glycan linked to residue N154. To analyze the replication of GUINEA-18, Vero E6 cells were infected at multiplicity of infection (m.o.i) of 0.1 and progeny virus productions were examined at various times post-infection (p.i.). using a conventional plaque-forming assay (Fig. 2A). Infection with MR766^MC^ served as a control. In comparison with MR766^MC^, virus progeny production of GUINEA-18 was significantly reduced by 2 log at 24 h reaching up to 3 log at 48 h p.i.. Infectious virus titer of GUINEA-18 peaked at 6 log PFU.mL^-1^ at 72 h p.i.. At this time point of infection, extensive cell death occurred in Vero E6 cells infected by MR766^MC^ but not GUINEA-18. The reduced progeny production of GUINEA-18 was associated to a significant delay in production level of E protein (Fig. 2B). At 48 h p.i., there was a 30-fold reduction in intracellular viral RNA (vRNA) production in Vero E6 cells infected with GUINEA-18 as compared with MR766^MC^ (Fig. 2C). At higher m.o.i of infection, GUINEA-18 progeny production was still reduced by 2 log at 48 h p.i. and this was associated to a low level of E protein expression as compared with MR766^MC^ (Fig. S2). A lactate dehydrogenase (LDH) assay was used as indicator of cellular damages in Vero E6 cells infected with ZIKV. Viability of Vero E6 cells infected with GUINEA-18 was essentially preserved till 72h p.i. whereas MR766^MC^ caused extensive cell death (Fig. 2D). Our data indicated that GUINEA-18 was a slowly replicating ZIKV in Vero E6 cells in comparison with MR766^MC^. Infection with GUINEA-18 does not produce major cytopathic effects in Vero E6 cells.

**Figure 2.**
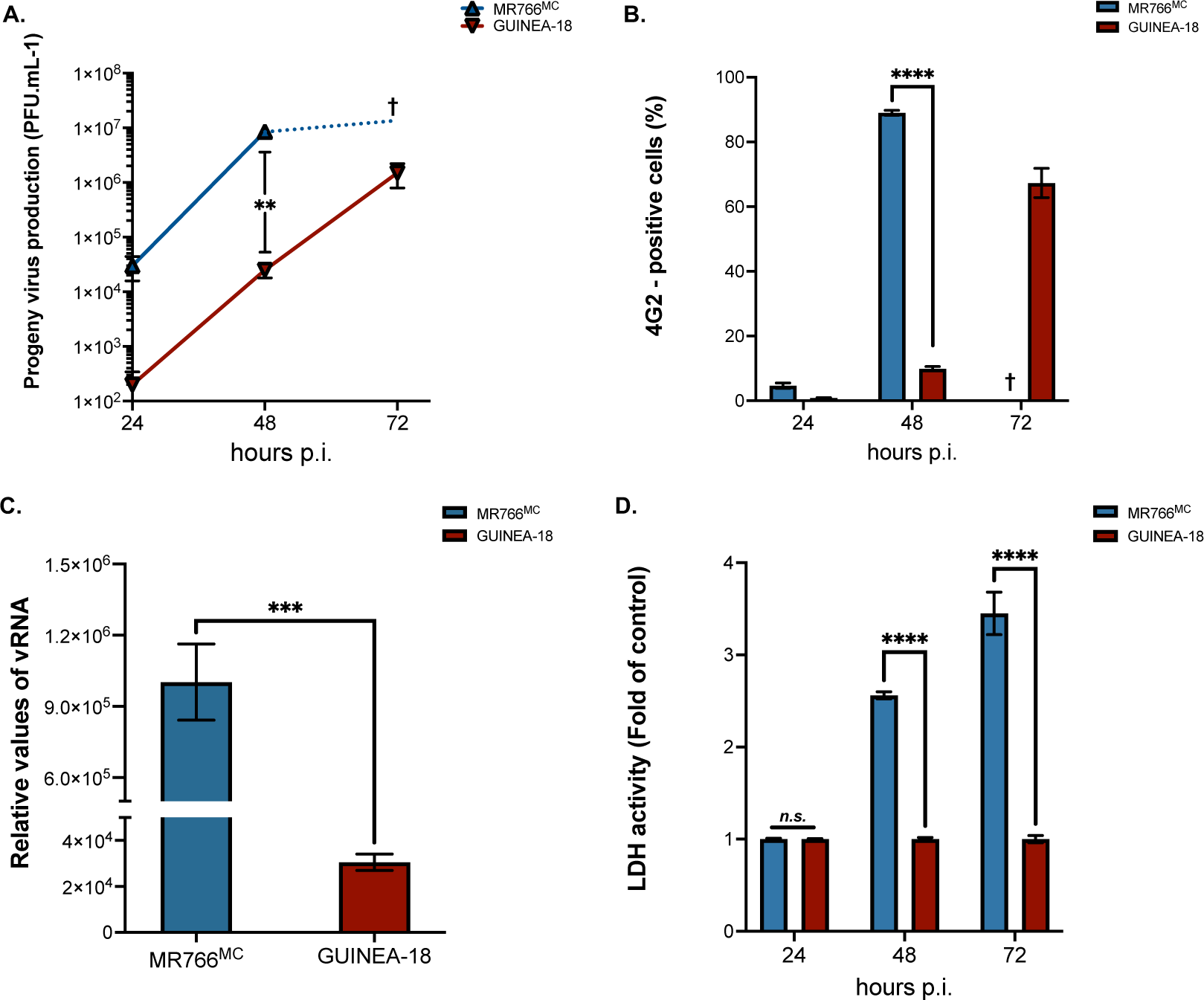
GUINEA-18 replication in non-human fibroblastic Vero E6 cells. Green monkey kidney fibroblastic Vero E6 cells were infected with MR766^MC^ and GUINEA-18 at m.o.i of 0.1 (**A** to **C**) or 1 PFU/cell (**D**). *In* (**A**), virus progeny productions at 24, 48 and 72 h p.i. were quantified by a standard plaque-forming assay. The cross signified that cell death occurred extensively during virus infection. *In* (**B**), FACS analysis was performed on ZIKV-infected cells using anti-*pan* orthoflavivirus E mAb 4G2 and the percentage of 4G2-positive cells was determined. *In* (**C**), intracellular viral RNA production was determined by RT-qPCR at 48 h p.i. RPLPO36B4 mRNA served as a house-keeping RNA control for normalization of samples. *In* (**D**), LDH activity was measured at 24, 48 and 72 h p.i. and expressed as a percentage relative to control. The results are the mean (± SEM) of three independent experiments. Asterisks indicate that the differences between experimental samples at each time point are statistically significant, using the unpaired *t* test (**** *P* < 0.0001, *** *P* < 0.001, ** *P* < 0.01; *n.s.*: not significant).

We examined the replication efficiency of GUINEA-18 in human pulmonary epithelial A549 cells and human microglia HCM3 cells that have been identified for their permissiveness to ZIKV infection. Firstly, A549 cells were infected with GUINEA-18 or MR766^MC^ at m.o.i. of 1. Analysis of viral growth identified a peak of progeny virus production reaching about of 7.5 log mL^-1^ in A549 cells infected for 72 h with GUINEA-18 (Fig. 3A). MR766^MC^ infection for 48 h was required to generate a similar infectious virus titer. At this timepoint of virus infection, GUINEA-18 virus progeny production was 10-fold lower as compared with MR766^MC^. There was associated to a 2-fold reduction in E protein expression level (Fig. 3B) and 8-fold reduction in vRNA production (Fig. 3C). Cytopathic effects were observed in MR766^MC^-infected A549 cells from 48 h p.i. whereas no loss of cell viability was observed with GUINEA-18 till 72 h p.i. (Fig. 3D). In contrast to MR766^MC^, GUINEA-18 replicated slowly in A549 cells and showed to cause little cytopathic effects. The slow replication of GUINEA-18 was also observed in HCM3 cells (Fig. S3). Taken together, these results showed that GUINEA-18 replicates slowly in different cell lines of various origin. The replication capacity of GUINEA-18 was reduced as compared with MR766^MC^.

**Figure 3.**
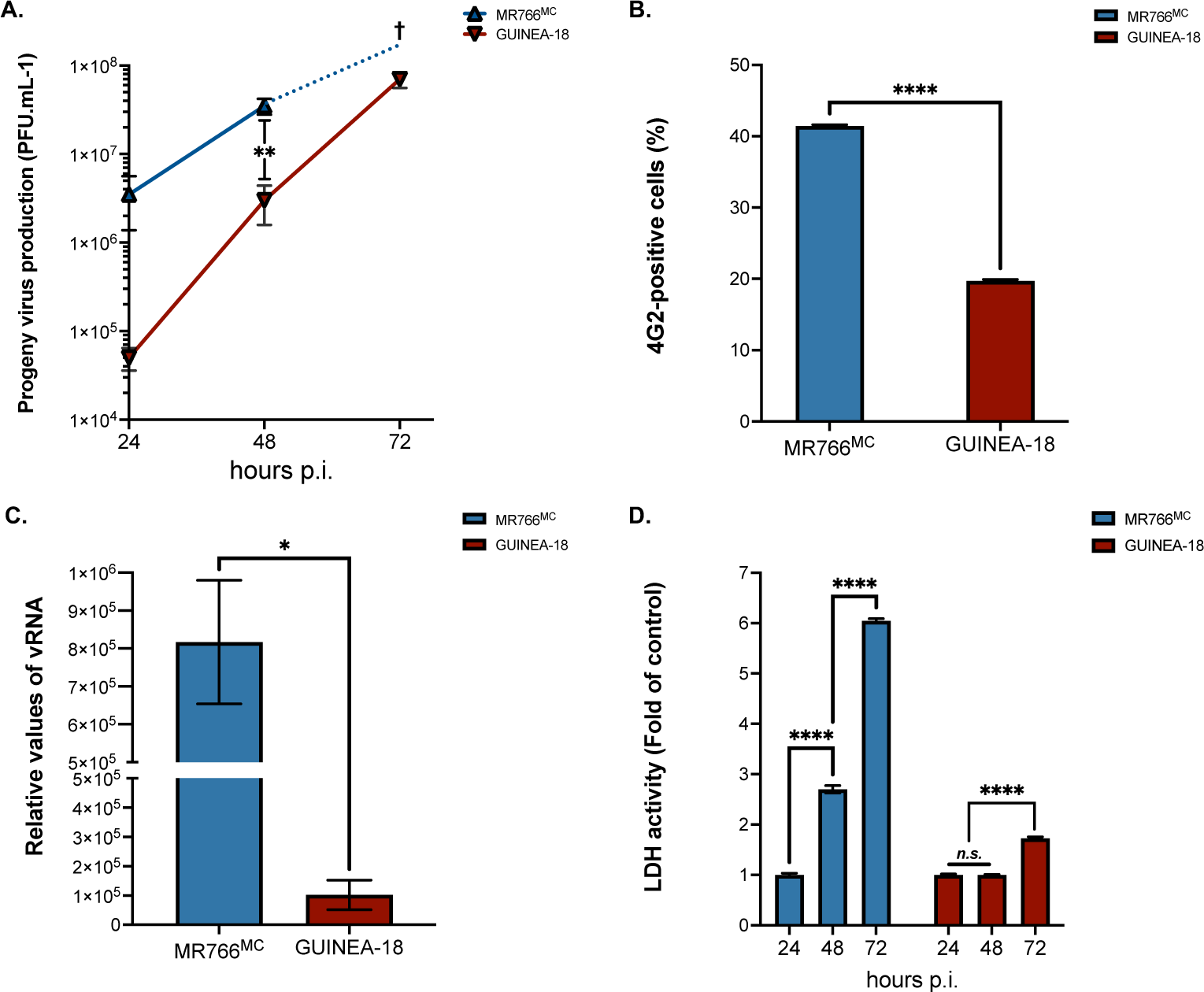
GUINEA-18 replication in human epithelial A549 cells. A549 cells were infected with MR766^MC^ and GUINEA-18 at m.o.i of 1. *In* (**A**), viral progeny production was determined at various times post-infection. The cross signified that cell death occurred extensively during virus infection. *In* (**B**), percentages of ZIKV-infected cells at 48 h p.i. were determined by FACS analysis using anti-E mAb 4G2 as primary antibody. *In* (**C**), intracellular viral RNA production was determined by RT-qPCR at 48 h p.i. RPLPO36B4 mRNA served as a house-keeping RNA control for normalization of samples. *In* (**D**), LDH activity was measured at 24, 48 and 72 h p.i. and expressed as a percentage relative to mock-infected cells (control). The results are the mean (± SEM) of three independent experiments. Asterisks indicate that the differences between experimental samples at each time point are statistically significant, using the unpaired *t* test (**** *P* < 0.0001, ** *P* < 0.01, ** *P* < 0.05; *n.s.*: not significant).

We assessed whether the lower replication capacity of GUINEA-18 in A549 cells was associated to a change in activation of antiviral innate immune responses as compared with MR766^MC^. ZIKV is targeted by IFN-type I such as IFN-β and antiviral effector proteins encoded by Interferon-Stimulated Genes (ISGs) through recognition of viral nucleic acids and molecular features associated with viral infection in the host-cell (22–25). The relative abundance expression of ISGs and IFN-β mRNA was assessed by RT-qPCR on total RNA isolated from ZIKV-infected A549 cells at 48 h p.i. (Fig. 4). As evidenced by analysis of fourteen representative ISGs, the ISG mRNA levels were significantly lower in A549 cells infected by GUINEA-18 in comparison with MR766^MC^. As a control of ISG protein production, ISG15 was weakly detected in A549 cells infected by GUINEA-18 compared with MR766^MC^ (Suppl. S4). The mRNA levels of IFN-β was also much lower in A549 cells infected by GUINEA-18 in comparison with MR766^MC^. Thus, infection of A549 cells with GUINEA-18 induces a lower ISG and IFN-β responses as compared with MR766^MC^.

**Figure 4.**
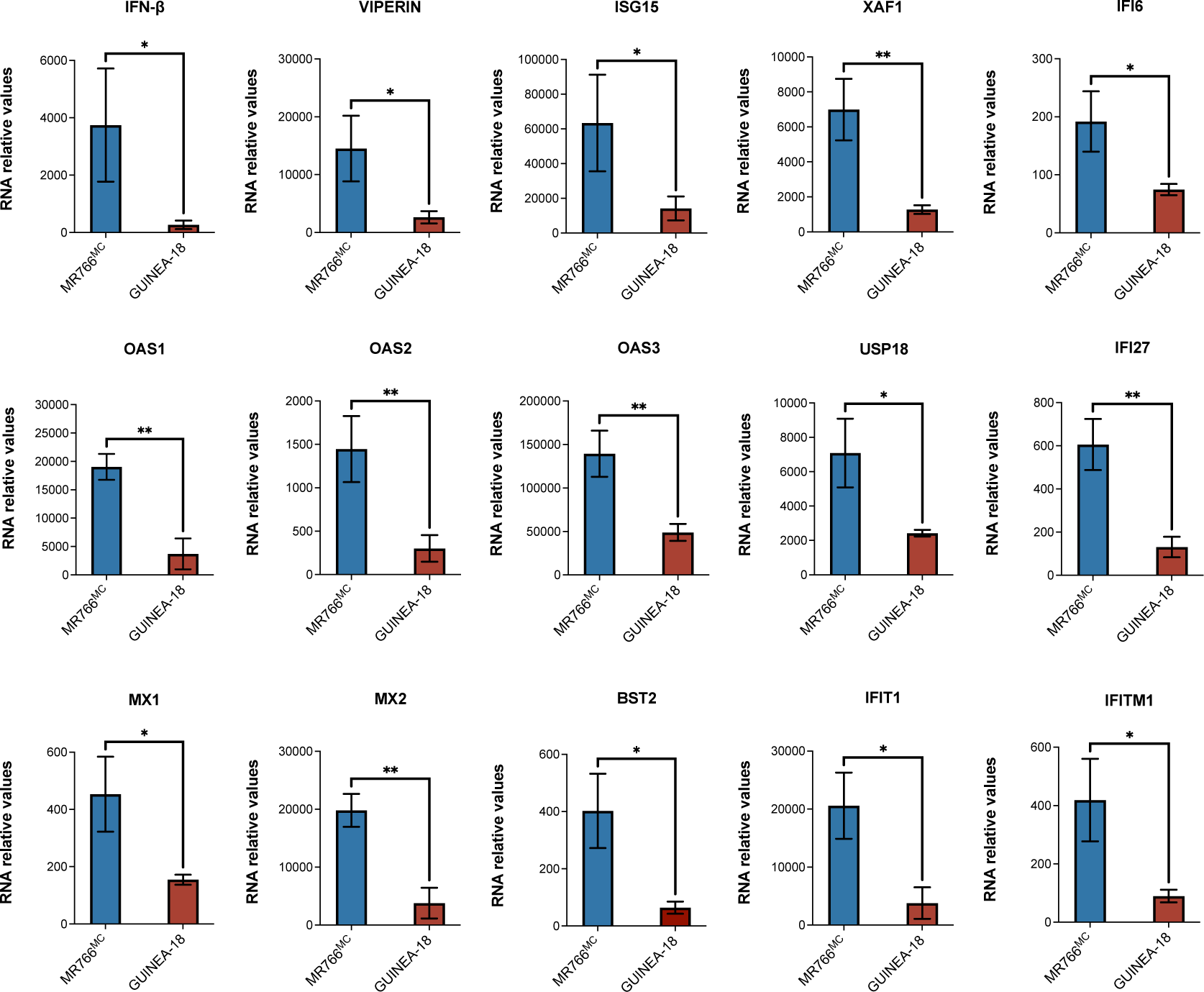
Expression of ISG and IFN-β mRNA in A549 cells infected by ZIKV. A549 cells were infected with GUINEA-18 or MR766^MC^ at m.o.i. of 1. The relative abundance of IFN-β and ISG mRNA was determined at 48 h p.i. by RT-qPCR. RPLPO36B4 mRNA served as a house-keeping RNA control for normalization of samples. Results are expressed as the fold-induction of IFN-β and ISG mRNA in ZIKV-infected cells relative to those in mock-infected cells. The results are the mean (± SEM) of three independent experiments. Asterisks indicate that the differences between experimental samples at each time point are statistically significant, using the unpaired *t* test (** *P* < 0.01, * *P* < 0.05).

### GUINEA-18 replication depends on NS1/NS4B proteins

We wondered whether a specific part of genomic viral RNA might play a role in attenuated replication of GUINEA-18 *in vitro*. The GUINEA-18 structural proteins with the N-glycosylated E protein may have an effect on virus binding efficiency in decreasing the host-cell susceptibility to viral infection (26,27). The importance of 5’ region of GUINEA-18 in viral attenuation was examined using GUINEA-18/MR766^5’region^ chimera in which the 5’NCR followed by the sequence coding for C, prM and E proteins of GUINEA-18 were replaced by the counterpart region of MR766^MC^ using genetic reverse experiments based on ISA method (Fig. 1B). There are two non-conserved nucleotides changes G19A and C41T in the 5’NCR and 8 amino acid substitutions in the structural protein region with 2 changes in C protein and 6 in E protein (Table 1). A549 cells were infected with GUINEA-18/MR766^5’region^ chimera or GUINEA-18 at m.o.i. of 1 (Fig. S5). Infection with MR766^MC^ served as a control. At 48 h p.i., there was no significant change in progeny virus production between GUINEA-18/MR766^5’region^ chimera and parental virus (Fig. S5A). This was also a similar expression level of E protein (Fig. S5B). These results rule out a major role for 5’NCR and the structural proteins in the lower replication efficacy of GUINEA-18 in comparison with MR766^MC^.

Comparative sequence analysis between GUINEA-18 and MR766^MC^ identified 8 mutations in the 3’NCR. The most notable 3’NCR mutations have been identified at positions 10508 in the stem loop II and 10635/10637 in a “dumbbell-like” structure. To determine whether the 3’NCR of GUINEA-18 might play a role in attenuated replication of GUINEA-18 *in vitro*, site-directed mutagenesis was conducted on the 3’NCR of MR766^MC^ in order to introduce the eight mutations from GUINEA-18 (Fig. 1B). The resulting MR766/3’NCR^GUINEA-18^ chimera and MR766^MC^ were compared for their replication in Vero E6 cells (Fig. S6). At 48 h p.i., the progeny virus production was comparable between MR766/3’NCR^GUINEA-18^ and the parental virus (Fig. S6A). This was associated to a similar percentage of ZIKV-infected cells (Fig. S6B). The MR766/3’NCR^GUINEA-18^ chimera was more cytopathic for Vero E6 cells than MR766^MC^ (Fig. S6C). Thus, it seems unlikely that 3’NCR impacts on the replication capability of GUINEA-18. Taken together, these results did not support a role for the structural protein region and the NCRs in attenuated replication of GUINEA-18 *in vitro*.

The above-mentioned results highlighted a possible role for the non-structural protein region in replication efficiency of GUINEA-18. Consequently, a first chimeric virus has been generated for which the MR766^MC^ coding region for NS1 to NS4B proteins was replaced by the counterpart from GUINEA-18 by ISA method (Fig. 1C). To facilitate the production of a such chimera, the NS5-N11S mutation was introduced into MR766^MC^ backbone. The replication of MR766/GUINEA^[NS1-NS4B]^ chimera was compared with MR766^MC^ in Vero E6 cells infected for 48 h at m.o.i. of 0.1 (Fig. 5). Infection with GUINEA-18 served as a control. Analysis of viral growth showed that insertion of GUINEA-18 NS1 to NS4B into MR766^MC^ strongly reduced virus progeny production (Fig. 5A). The curves of viral multiplication were comparable between MR766/GUINEA-18^[NS1-NS4B]^ chimera and GUINEA-18. The two viruses were indistinguishable for the E protein expression level (Fig. 5B) and intracellular vRNA rate in Vero E6 cells (Fig. 5C). Late in infection, MR766/GUINEA-18^[NS1-NS4B]^ preserved cell viability as it has been observed with GUINEA-18 (Fig. 5D). These results showed that insertion of GUINEA-18 NS1/NS4B proteins in MR766^MC^ was efficient to affect virus replication capability.

**Figure 5.**
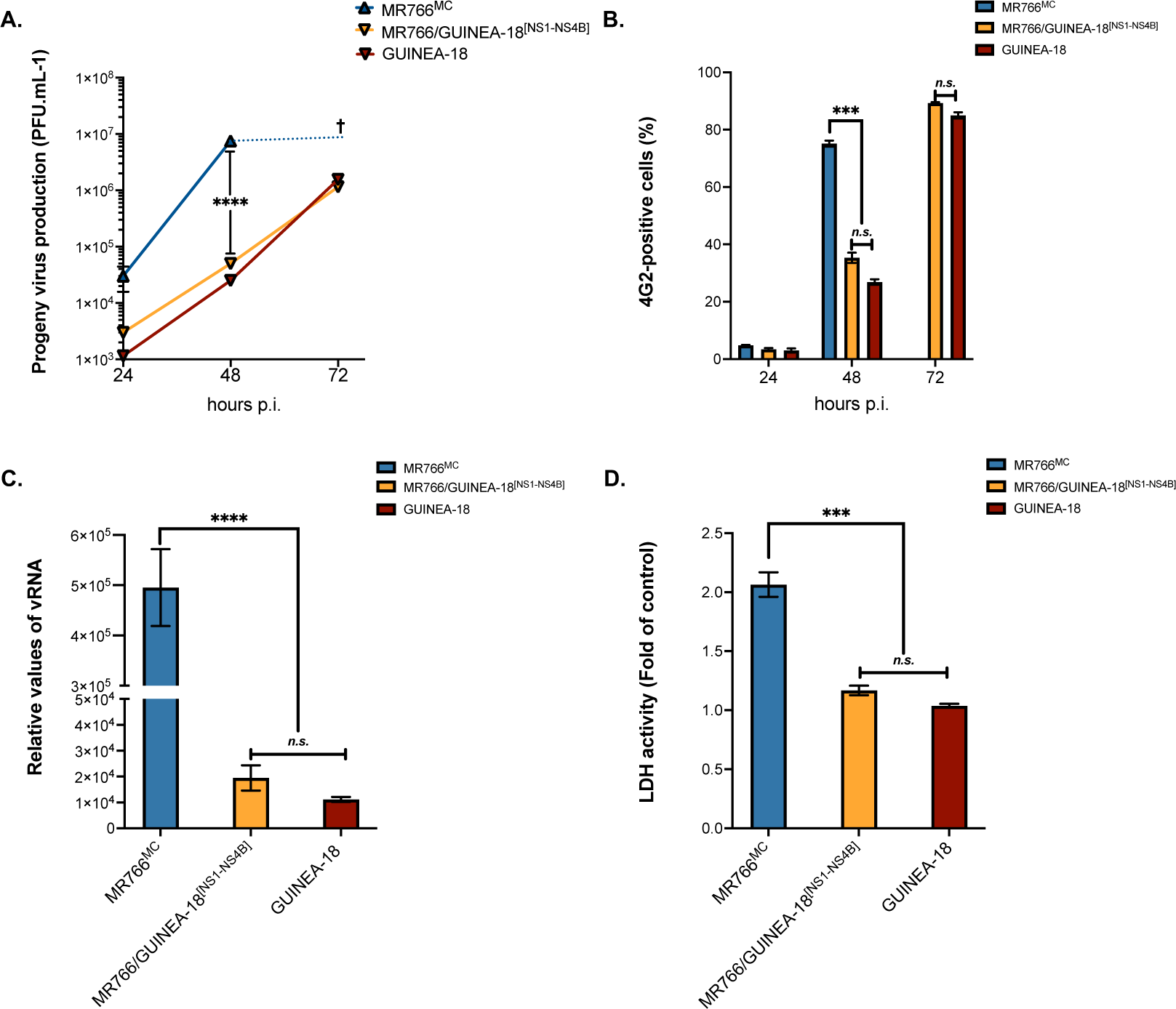
MR766/GUINEA-18^[NS1-NS4B]^ replication in Vero cells. Vero E6 cells were infected with MR766/GUINEA-18^[NS1-NS4B]^ chimeric virus or parental viruses at m.o.i of 0.1. *In* (**A**), virus progeny production at 24h, 48h, and 72h. *In* (**B**), percentage of ZIKV-infected cells at 24h, 48h, and 72h measured by FACS analysis using mAb 4G2. *In* (**C**), RT-qPCR was performed on viral RNA (vRNA) extracted from Vero E6 cells infected for 48h with ZIKV. RPLPO36B4 mRNA served as a house-keeping RNA control for normalization of samples. *In* (**D**), LDH activity was measured at 72 h p.i. The results are the mean (± SEM) of three independent experiments. Asterisks indicate that the differences between experimental samples at each time point are statistically significant, using the unpaired *t* test or ANOVA (**** *P* < 0.0001, *** *P* < 0.001; *n.s*.: not significant).

We next evaluated the replication capability of MR766/GUINEA-18^[NS1-NS4B]^ chimera in A549 cells (Fig. 6). Infection of A549 cells with the chimera resulted in a significant reduction in virus progeny production associated to a lower production of intracellular vRNA and E protein expression at 48 h p.i. as compared with MR766^MC^ (Fig. 6). MR766/GUINEA-18^[NS1-NS4B]^ chimera and GUINEA-18 were indistinguishable reinforcing the notion that GUINEA-18 NS1/NS4B proteins affect virus replication capability in different cell lines.

**Figure 6.**
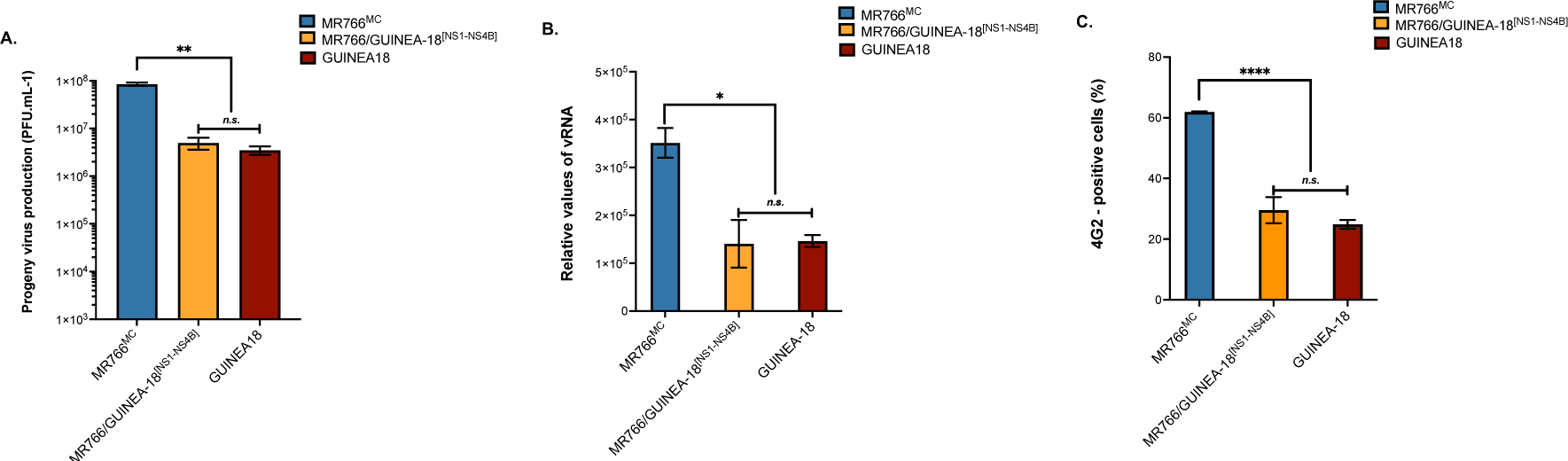
MR766/GUINEA-18^[NS1-NS4B]^ replication in A549 cells. A549 cells were infected for 48h with MR766/GUINEA-18^[NS1-NS4B]^ chimera or parental viruses at m.o.i of 1. *In* (**A**), virus progeny production. *In* (***B***), RT-qPCR was performed on viral RNA (vRNA) extracted from ZIKV-infected Vero E6 cell. RPLPO36B4 mRNA served as a house-keeping RNA control for normalization of samples. *In* (**C**), percentage of ZIKV-infected cells measured by FACS analysis using mAb 4G2. The results are the mean (± SEM) of three independent. Asterisks indicate that the differences between experimental samples at each time point are statistically significant, using the unpaired *t* test or ANOVA (**** *P* < 0.0001, ** *P* < 0.01; * *P* < 0.05; *n.s*.: not significant).

To determine whether the features of GUINEA-18 depend on all or some NS proteins, we produced the additional chimeric viruses, MR766/GUINEA-18^[NS1-NS3]^ and MR766/GUINEA-18^[NS1-N2A]^, containing the GUINEA-18 NS1/NS2AB/NS3 or NS1/NS2A genes into MR766^MC^ backbone, respectively (Fig. 1C). The replication capability of chimeric viruses was analyzed in Vero E6 cells infected at the m.o.i. of 0.1 and compared with parental viral clones (Fig. 7A). Infection with MR766/GUINEA^[NS1-NS4B]^ chimera served as a control. Analysis of virus progeny production revealed that MR766^MC^ chimeric viruses containing GUINEA-18 NS1/NS3 or NS1/NS2A proteins have little or no effect on virus replication (Fig. 7A). This was associated to a comparable loss of cell viability between chimeric viruses (Fig. 7B). Since NS4A protein is strictly conserved between GUINEA-18 and MR766^MC^, the lack of effect of GUINEA-18 NS1/NS3 proteins on MR766^MC^ replication emphasizes the major role of NS4B in the attenuated phenotype of GUINEA-18.

**Figure 7.**
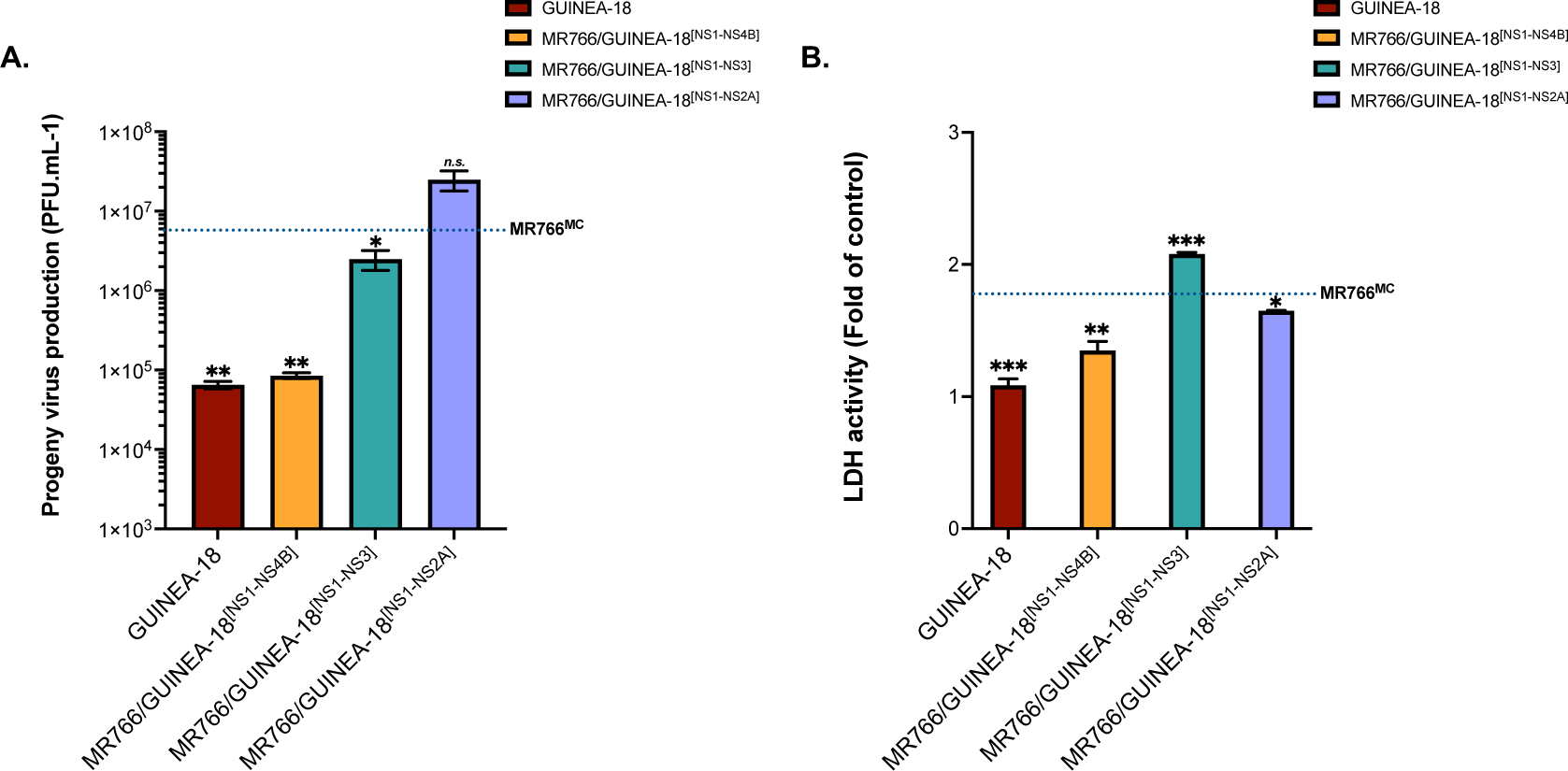
Impact of GUINEA-18 NS proteins on MR766^MC^ replication. Vero E6 cells were infected with MR766/GUINEA-18 chimeric viruses containing GUINEA-18 NS proteins or parental ZIKV at m.o.i. of 0.1. *In* (**A**), virus progeny production at 48h p.i. The dotted line indicates MR766^MC^ progeny production. *In* (**B**), LDH activity was measured at 72 h p.i. The dotted line indicates the rate of LDH release with MR766^MC^. The results are the mean (± SEM) of three independent experiments. Statistical analysis relative to MR766^MC^ values was noted. Asterisks indicate that the differences between experimental samples at each time point are statistically significant, using the unpaired *t* test (*** *P* < 0.001, ** *P* < 0.01, * *P* < 0.05; *n.s*.: not significant).

The above results suggest that replication capability of GUINEA-18 depends on NS4B protein. Among seven amino-acid substitutions that differentiate NS4B of ZIKV-1555 and MR766, four have been identified at positions 11, 20, 24 and 26 (Table 1). The ZIKV-15555 residues R20 and T24 have been also identified in West African ZIKV strains Senegal-Kedougou 2011 and Senegal-Kedougou 2015 (European Nucleotide Archive accession number n°PRJEB39677) that have been isolated from mosquito pools in Casamance region of Senegal in 2011 and 2015, respectively (15). The residues R20 and T24 were not observed in Asian and Asian-related American viral strains. A change from non-polar amino-acid alanine to polar residue threonine at position 24 might have an effect on NS4B conformation. A three-dimensional structure of the N-terminal region of NS4B was performed by modelling on Phyre^2^ (28) allowing *de novo* peptide structure prediction (Fig. S7). Structural analysis showed that ZIKV-1555 NS4B residues 1 to 16 and 35 to 50 have propensity for forming a-helical structure and transmembrane helix, respectively. The NS4B amino-acid substitutions at positions 20, 24, and 26 that differentiate ZIKV-15555 from MR766 have been identified in a non-ordered structure between helix a1 and TM1 (29).

We investigated whether the NS4B mutations that differentiate ZIKV-15555 from MR766 influence protein expression. Recombinant 2K-NS4B (hereafter intitled rNS4B) proteins derived from ZIKV-15555, MR766, and BeH819015 have been expressed in A549 cells using a vector plasmid pcDNA3. The rNS4B proteins were C-terminally tagged with a FLAG peptide. The ZIKV-15555 rNS4B protein expressed without the 2K peptide served as control. The rNS4B protein expression in transfected A549 cells was verified by FACS analysis using anti-FLAG antibody (Fig. S8). The levels of rNS4B protein expression were comparable among the different rNS4B proteins expressed in A549 cells (Fig. 8). Immunoblot assays using anti-FLAG antibody allowed the detection of MR766 and ZIKV-15555 rNS4B proteins in A549 cells (Fig. 8A) and HCM3 cells (Fig. 8B). Their profile migration was comparable with an apparent molecular weight similar to rNS4B mutant without 2K indicating that the N-terminal peptide was correctly processed from the N-terminus of protein. The change in profile migration of BeH819015 NS4B protein as compared with ZIKV-15555 and MR766 could be due to specific residues that have been identified in Asian and Asian-related American viral strains. We can conclude that the seven NS4B amino-acid substitutions that differentiate ZIKV-15555 from MR766 have no obvious effect on protein expression.

**Figure 8.**
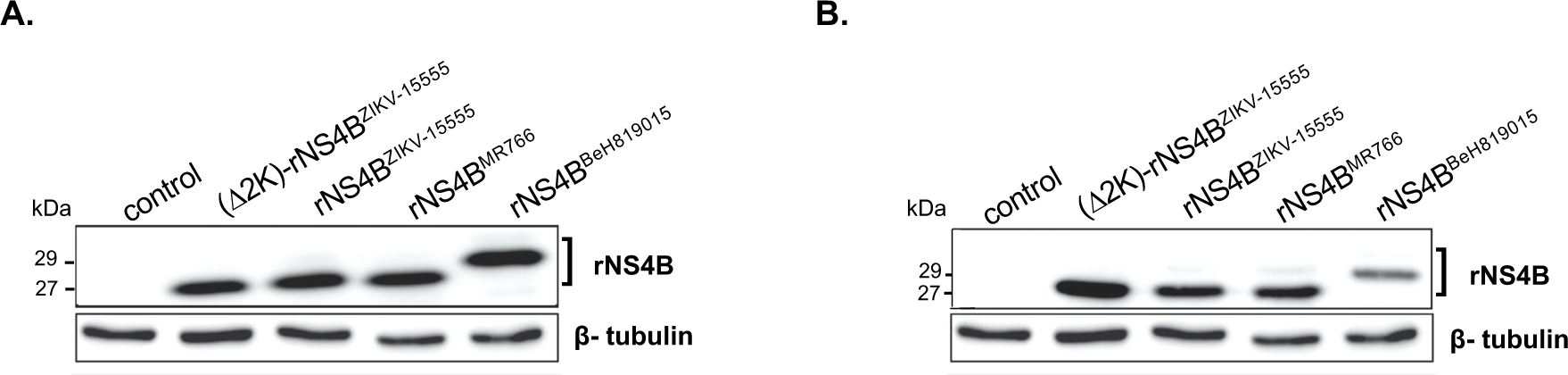
Intracellular expression of recombinant ZIKV NS4B protein. A549 (**A**) and HCM3 (**B**) cells were transfected for 24 h with pcDNA3 expressing rNS4B protein from ZIKV-15555 (rNS4B^ZIKV-15555^), MR766 (rNS4B^MR766^), or BeH819015 (rNS4B^BeH819015^). The rNS4B^ZIK-15555^ protein mutant lacking the 2K peptide is indicated as (Δ2K)-rNS4B^ZIK-15555^. Cell lysates in RIPA buffer were assessed by immunoblot assays with anti-epitope antibody FLAG as the primary antibody. The β-tubulin protein served as a protein-loading control. The positions of rNS4B are indicated.

To better understand the role of NS4B in attenuation replication of GUINEA-18, we produced MR766^MC^/NS4B^GUINEA-18^ chimera (Fig. 1) and MR766^MC^-NS4B mutant containing the GUINEA-18 NS4B protein or its the four N-terminal residues S11/R20/T24/I26 including V180, respectively. Vero E6 cells were infected with MR766^MC^/GUINEA^NS4B^ chimera and MR766^MC^-NS4B mutant at m.o.i. of 0.1 (Fig. 9). Parental viruses and MR766/GUINEA-18^[NS1-NS4B]^ chimera served as controls. Analysis of MR766^MC^/NS4B^GUINEA-18^ revealed that insertion of GUINEA-18 NS4B in MR766^MC^ caused no reduction in virus progeny production (Fig. 9A) and expression level of E protein (Fig. 9B). By contrast, infection with MR766^MC^-NS4B^(S11/R20/T24/I26/V180)^ mutant resulted in low virus progeny production comparable to that observed with GUINEA-18 and MR766/GUINEA-18^[NS1-NS4B]^ (Fig. 9C). This was associated to weak expression level of E protein (Fig. 9D). Also, the viability of Vero E6 cells infected with MR766^MC^-NS4B mutant was preserved at 72 h p.i. (Fig. 9E). However, the effects of NS4B mutations on MR766^MC^ replication were not observed in A549 cells raising the possibility that their impact on virus replication depends on interactions of viral protein with cellular factors in a cell-type-dependent manner (Fig. 9F). These results suggested that the N-terminal residues of GUINEA-18 NS4B influence virus replication capability but their impact may depend on environment.

**Figure 9.**
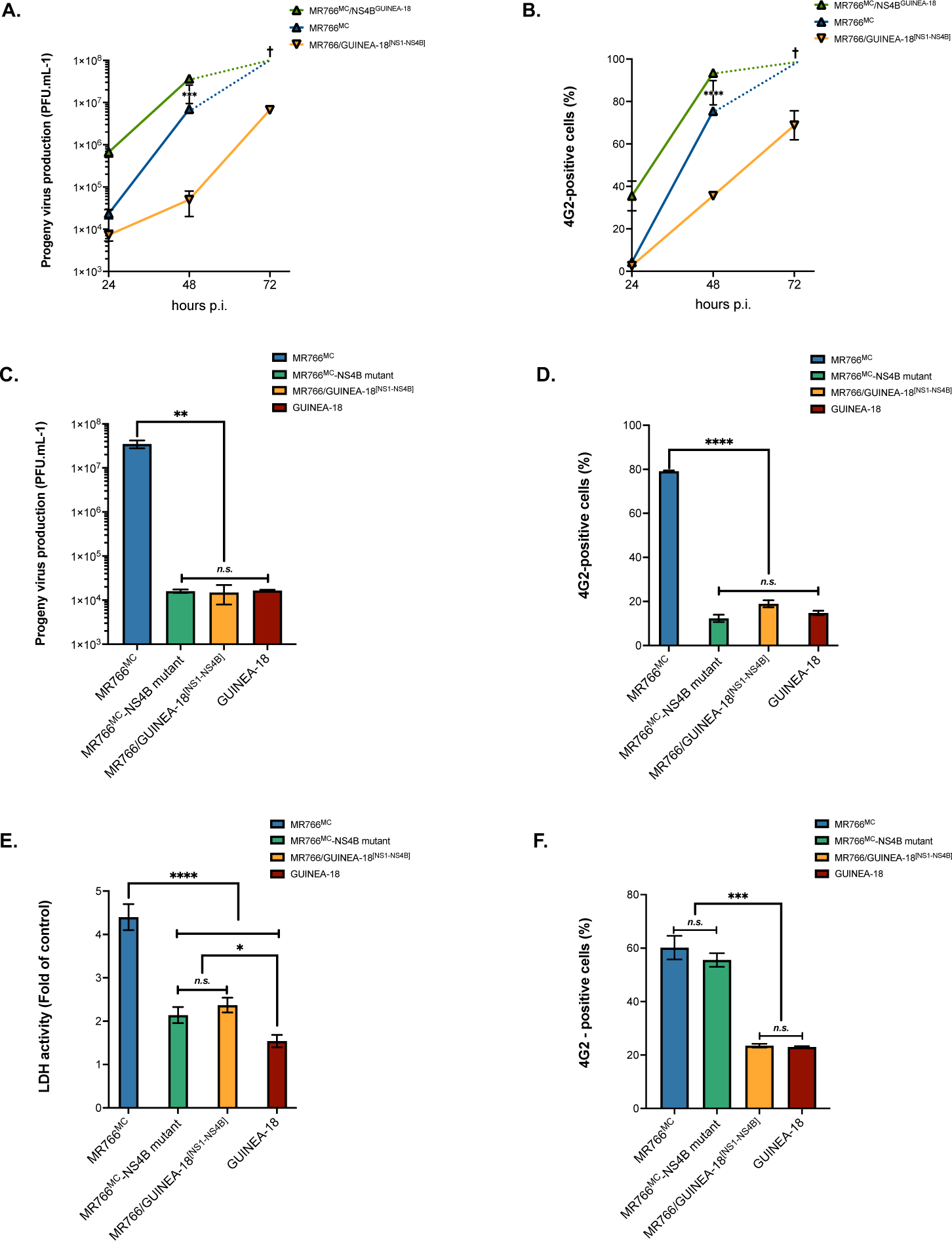
Effect of GUINEA-18 NS4B on MR766^MC^ replication. Vero E6 cells (**A-E**) and A549 cells (**F**) were infected with GUINEA-18, MR766^M^, MR766^MC^ mutant containing the GUINEA-18 NS4B residues S11/R20/T24/I26/V180 (MR766^MC^-NS4B mutant) or MR766^MC^ chimeric virus containing the GUINEA-18 NS4B protein (MR766^MC^/NS4B^GUINEA-18^). MR766/GUINEA-18^[NS1-NS4B]^ served as a control. *In* (**A-B**), Vero E6 cells were infected with MR766^MC^/NS4B^GUINEA-18^ at m.o.i 0.1. Virus progeny production (**A**), and percentage of ZIKV-infected cells measured by FACS analysis using mAb 4G2 (**B**). The cross signified that cell death occurred extensively during virus infection. *In* (**C-E**), Vero E6 cells were infected with MR766^MC^-NS4B mutant at m.o.i 0.1. Virus progeny production (**C**), and percentage of ZIKV-infected cells measured by FACS analysis using mAb 4G2 (**D**) were determined at 48 h p.i. LDH activity was measured at 72 h p.i. **(E).** *In* (**F**), A549 cells were infected with MR766^MC^-NS4B mutant at m.o.i 1. The percentages of ZIKV-infected cells measured by FACS analysis using mAb 4G2 were determined at 48 h p.i. The results are the mean (± SEM) of two or three independent experiments. Asterisks indicate that the differences between experimental samples at each time point are statistically significant, using the unpaired *t* test or ANOVA (**** *P* < 0.0001, *** *P* < 0.001, ** *P* < 0.01; * *P* < 0.05; *n.s*.: not significant).

### GUINEA-18 NS1/NS4B proteins prevent stress granule assembly

In response to RNA virus infection, cellular stress granules (SGs) are assembled as cytoplasmic condensates which can sequester viral RNA and proteins restricting viral growth (30–32). Orthoflaviviruses including ZIKV have developed strategies to prevent SG formation facilitating viral replication and also limiting antiviral signaling activation (32–34). We wondered whether GUINEA-18 NS1/NS4B proteins may have an effect on SG formation. To assess SG assembly in A549 cells, eGFP reported was fused in-frame to the N-terminus of RasGAP SH3 domain-binding protein (G3BP) which is considered as a robust SG marker (33,35,36). The sugar sorbitol as a physiological osmotic and oxidative stressor, was used to drive GFP-G3BP fusion protein to SGs (37,38). Confocal fluorescence microscopy identified eGFP-positive condensates in A549 cells incubated with 0.4M sorbitol for 1.5h with an average surface area of 2.5 µm^2^ (Fig. S9).

To evaluate the effect of GUINEA-18 NS proteins on SG assembly, A549 cells were transfected with plasmid expressing eGFP-G3BP fusion protein for 6 h and then infected for 40 h with MR766/GUINEA^[NS1-NS4B]^ chimera at m.o.i. of 2 (Fig. 10). Both MR766^MC^ and GUINEA-18 served as controls of ZIKV-specific SG formation blockade. The ZIKV-infected cells were visualized by IF assay using anti-E mAb 4G2. Analysis of A549 cells infected for 40 h with MR766^MC^ and then stressed with 0.4M sorbitol for 1.5 h identified large eGFP-positive condensates in the cytoplasm (Fig. 10A). Almost fifty 4G2-positive cells were scored for the number of eGFP-positive condensates per cell profile. An average of seven defined granules per cell was observed in A549 cells infected by MR766^MC^ (Fig. 10B). Most of the eGFP-positive condensates had a surface area ranging from 3 to 9 µm^2^ with an average of approximately 4.1 µm^2^ (Fig. 10C). In contrast, A549 cells infected with GUINEA-18 and stressed with sorbitol were essentially free of eGFP-positive condensates in the cytoplasm (Fig. 10B). Thus, GUINEA-18 is greatly efficient to impair SG formation under environmental stress. This was a smaller number of defined granules (average of 3 per cell) in A549 cells infected by MR766/GUINEA^[NS1-NS4B]^ chimera as compared with parental virus (Fig. 10B). The number of eGFP-positive condensates with a surface area ranging from 3 to 9 µm^2^ was reduced by nearly 50% (Fig. 10C). The surface area average of defined granules was approximately 2.5 µm^2^ (Fig. 10C). Thus, insertion of GUINEA-18 NS1/NS4B proteins in MR766^MC^ resulted in a significant decrease in the number and size of SGs in A549 cells stressed with as an environmental stressor. Together, these results showed that GUINEA-18 has a greater capacity to affect SG assembly as compared with MR766^MC^. The NS1/NS4B proteins have been identified as playing a key role in the capacity of GUINEA-18 to prevent SG assembly in response to ZIKV infection.

**Figure 10.**
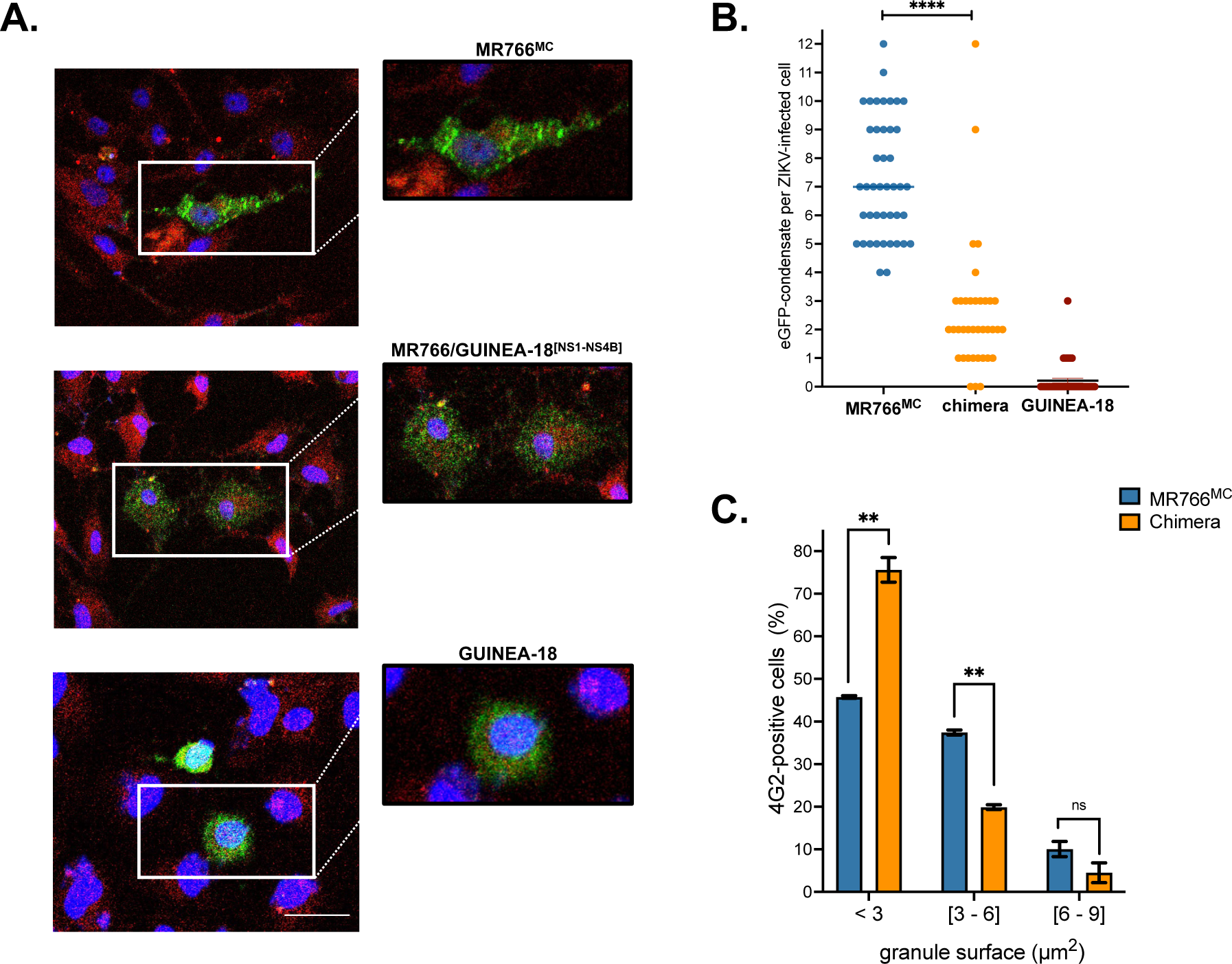
Stress granule formation in ZIKV-infected cells stressed with sorbitol. A549 cells were transfected with pcDNA3/eGFP-G3BP for 6 h and then infected with ZIKV at m.o.i. of 2. At 40 h p.i., cells were stressed with 0.4 M sorbitol for 1.5 h. *In* (**A**), cells infected with MR766^MC^, GUINEA-18, or MR766/GUINEA-18^[NS1-NS4B]^ chimera were identified using anti-E mAb 4G2. Cells positive for ZIKV E (red) and eGFP-G3BP (green) proteins were examined by confocal microscopy analysis. Nuclei were stained with DAPI (blue). *In* (**B**), cells positive for E protein expression (n ∼ 50) were scored for the number of eGFP-positive condensates per ZIKV-infected cell. Values from 4 independent experiments are presented. Asterisks indicate that the differences between experimental samples at each time point are statistically significant, using 2-way ANOVA test (**** *P* < 0.0001). *In* (**C**), cells positive for E protein and eGFP-G3BP expression (n ∼ 100) were scored for the surface of EGFP-positive condensates as small (< 3 µm^2^), medium (3 - 6 µm^2^) and large (6 - 9 µm^2^) defined granules. Values from two independent experiments are presented. Asterisks indicate that the differences between experimental samples at each time point are statistically significant, using the unpaired *t* test (\*\**P* < 0.01; *n.s*.: not significant). Scale bar, 50 μm.

## DISCUSSION

Most of studies having for objectives to understand the transmission, biology, and pathogenicity of emerging ZIKV are still carried out using epidemic strains of Asian lineage. Too little information is available on the replication properties of contemporary African ZIKV strains despite their high of transmission level by *Aedes* mosquito vectors and their teratogenic potential for humans. It has been proposed that the greater level of transmissibility and pathogenicity of African ZIKV strains might be the cause of fetal loss rather than neurodevelopmental malformations (15). This could explain the difficulties to estimate the adverse pregnancy outcomes associated to ZIKV infection in sub-Saharan Africa (39–41).

It is of priority to improve our knowledge on features of recently isolated ZIKV strains from West Africa. Genomic RNA of viral strain ZIKV-15555 that has been sequenced from an individual who had been exposed to ZIKV in Guinee (Faranha region) in 2018 has been considered as suitable for studying the contemporary West Africa viral strains. Here, an infectious molecular clone GUINEA-2018 that has been obtained from ZIKV-15555 RNA sequence by reverse genetic approach based on the ISA method. Viral clone MR766^MC^ from historical African ZIKV strain MR766 has been chosen as prototype model of ZIKV of African lineage in our study (19). Viral clones GUINEA-18 and MR766^MC^ share a high degree of amino-acid homology (98.6% of identity) with only six amino-acid substitutions in the E protein in 70 years of distance. Also, there were only few nucleotide mutations between the 5’NCRs and 3’NCRs of the two viral clones. *In vitro* cell cultures studies of GUINEA-18 replication were conducted in comparison with MR766^MC^. Viral growth analysis showed that GUINEA-18 replication is markedly attenuated in Vero E6 cells as compared with MR766^MC^. The attenuated replication capability of GUINEA-18 coincided with a low vRNA production rate and reduced production of intracellular E protein. Cell viability was mostly preserved with GUINEA-18 whereas extensive cell death occurred with MR766^MC^ within the three days of infection. The replication capability of GUINEA-18 was also attenuated in A549 cells leading to a slight loss of cell viability. Infection with GUINEA-18 resulted to a moderate up-regulation of IFN-β and ISGs expression in A549 cells as compared with MR766^MC^. The weak induction of innate immune responses might be a direct consequence of reduced replication capability of GUINEA-18. It is also tempting to speculate that GUINEA-18 has evolved exquisite strategies to evade host innate immune defenses within the host cells that it infects.

In an effort to identify genetic determinants possibly involved in the lower replication capability of GUINEA-18, chimeric viruses have been generated by interchanging viral sequences from the two African ZIKV strains using the ISA method. Two mutations have been identified between the 5’ NCR of GUINEA-18 and MRC766^MC^. Noteworthy, their structural protein regions only differ by 2 and 6 amino acid substitutions in the C and E proteins, respectively. The prM protein which has been proposed as viral factor playing a key role in the neuropathogenicity of epidemic Asian ZIKV strains was unchanged between GUINEA-18 and MR766^MC^. GUINEA-18 E protein includes a flexible glycan loop (GL, residues E-145 to E-164) region where a glycan is linked to residue N154 (42). The N-glycosylation of the GL region has been also observed with epidemic ZIKV strains of Asian lineage (42). In contrast, MR766^MC^ includes the T156I substitution which abrogates the N-glycosylation site resulting to in a non-glycosylated E protein for historical African viral strain (42). The replacement of GUINEA-18 structural protein region by the counterpart from MR766^MC^ generated a chimeric virus showing similarity in replication and cytotoxicity to parental virus. This excludes a role for the C and E proteins in attenuation of GUINEA-18 *in vitro*. The permutation of MR766^MC^ 3’NCR by the counterpart from GUINEA-18 that differs by eight mutations had no effect on MR766^MC^ replication precluding a role for the 3’end of genomic RNA in the attenuated replication of GUINEA-18.

Forty-one amino-acid substitutions have been identified between the NS proteins of ZIKV-15555 and MR766 (Table 1). Only NS2B protein was unchanged between the two viral strains. Swapping the MR766^MC^ coding region for NS1 to NS4B proteins by the GUINEA-18 counterpart was shown to result in an attenuated chimeric virus exhibiting attenuated replication and restricted cytotoxicity to Vero E6 and A549 cells similar to GUINEA-18. Our data identified a central role for NS1 to NS4B proteins that compose VRCs in the replication capability of GUINEA-18. It is of note that the involvement of NS5 protein was not investigated in our study. Based on MR766 chimeric viruses, we found that GUINEA-18 NS4B plays an essential role in virus attenuation. From the seven NS4B amino-acid substitutions that differentiate GUINEA-18 from MR766^MC^, four have been identified in the N-terminal region of the protein with K20 and T24 residues as unique for contemporary West African ZIKV strains. We propose a role for the N-terminal residues of NS4B in attenuated replication of GUINEA-18 but their effects depend on cellular environment and presumably interactions with other NS proteins within the VRCs.

The formation of SGs as biomolecular condensates including G3BP protein is induced by various stresses including viral infections (31,35). Inhibition of SG formation or SG disassembly contributes to strategies by which RNA viruses can hijack innate immune responses to their own benefit (43). Consistent with the notion that orthoflaviviruses prevent SG assembly (32), ZIKV-specific SG formation blockade was observed in A549 cells regardless viral strain tested. This observation agreed with the proposal that ZIKV developed efficient strategies to suppress GS formation and/or trigger SG disassembly. In A549 cells infected with ZIKV and then stressed with sorbitol as environmental SG inducer, GUINEA-18 showed a great efficacy to prevent SG formation as compared with MR766^MC^. This raises the question of whether blockade of SG formation contributes the weak induction of ISGs and IFN-β in A549 cells infected by GUINEA-18. The sorbitol-induced SG formation of SGs was greatly affected in A549 cells infected with a MR766^MC^ chimera containing the GUINEA-18 NS1 to NS4B proteins emphasizing a potential role for NS proteins in ZIKV-specific SG formation blockade. It is still unknown which specific NS protein(s) may be responsible for suppressing SG formation. Whether the attenuated replication and restricted cytotoxicity of GUINEA-18 associated to a weak induction of innate immune responses are determined by the capacity of ZIKV to suppress SG formation in link with NS proteins is a critical issue that remains to be investigated.

In our study, molecular clone-based comparative analysis with historical African ZIKV strain MR766 allowed to identify contemporary African viral strain ZIKV-15555 as attenuated virus *in vitro*. Our data indicated that the attenuated phenotype of viral strain essentially depends on NS1 to NS4B proteins with a particular emphasis on N-terminal region of NS4B. What might be the specific role of each NS proteins on attenuated replication and restricted cytotoxicity of ZIKV-15555 *in vitro*? The thin regulation of NS protein interactions that is required for efficient VRCs where genome replication and transcription occur in organelle-like structures might be distinct for ZIKV-15555 in comparison to MR766. This would be consistent with the possibility that viral clone of ZIKV-15555 has a stronger efficacy than MR766 in suppressing SG formation or by facilitating SG disassembly. The weak induction of ISGs in the host-cells infected with viral clone of ZIKV-15555 could rely to the capacity of virus to prevent SG formation. It is therefore of priority to determine whether the NS proteins of contemporary African viral strains attribute to ZIKV a greater ability to hijack antiviral innate immunity by limiting SG assembly in the host-cells that they infect. This might play an important role in the virulence of contemporary Africa viral strain ZIKV-15555 infection that remains to be elucidated *in vivo* (44).

## Acknowledgements

We do want to thank L. Lambrechts for his input and insight into sequence data for ZIKV strains Senegal-Kedougou. We thank G. Gadea, N. Jouvenet, B. Mesmin, C. Atyame-Nten and PE. Ceccaldi for their interest in the study. We gratefully acknowledge J. Andries for help with virus productions. We thank PIMIT members for helpful discussions. We thank PICT platform (University of Reims Champagne-Ardenne) for imaging core facilities. This work was funded by the French government as part of France 2030 with the support of ANRS I MIE through the ANRS-23-PEPR-MIE 0004 project intitled CAZIKANO. D.M. was supported by a doctoral scholarship from the University of La Réunion (Ecole doctorale STS), funded by the French ministry MESRI.

## Author contributions

P.D. and D.M. conceived and designed the experiments. D.M. performed the experiments. P.D., M-P.C, A.K., D.M. analyzed the data. P.D. and C.E.K contributed to reagents/materials/analysis tools. P.D, and D.M. wrote the original draft preparation. D.M, M-P.C, C.K., A.K, P.D. wrote review and editing. All authors have read and agreed to the published version of the manuscript.

## Supplemental data (Machmouchi et al.)

**Figure S1.**
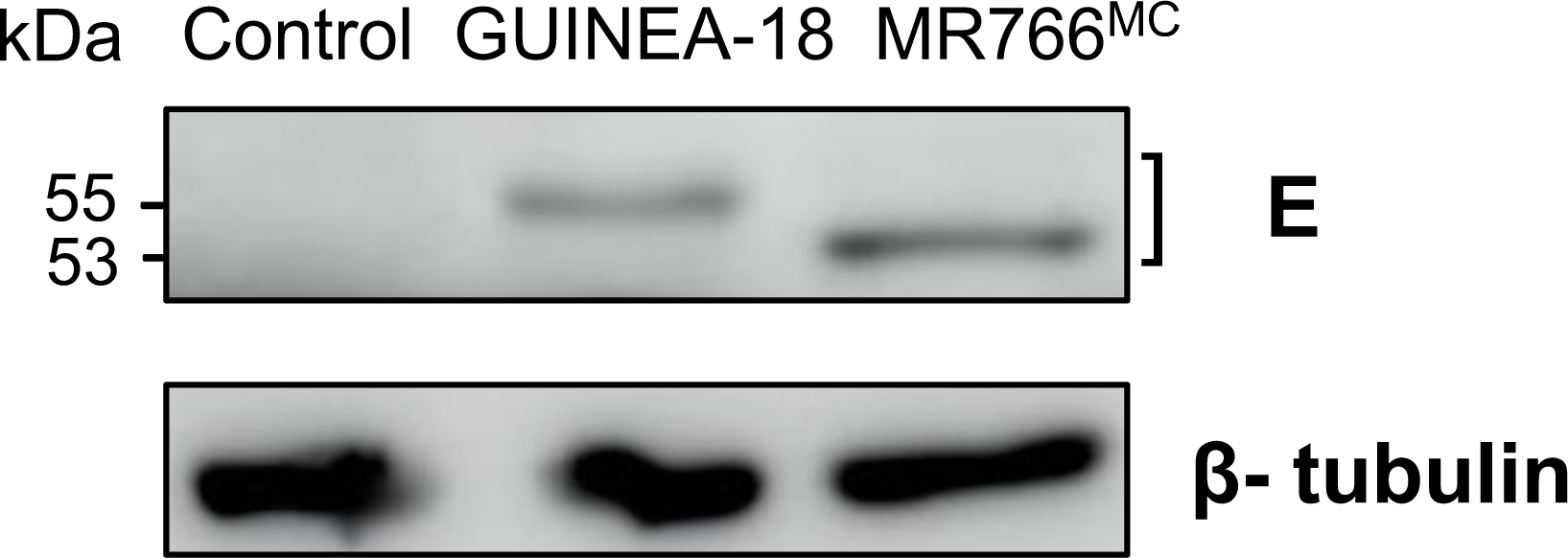
Glycosylation profile of ZIKV E protein. Vero cells were infected for 48 h with infectious molecular clones GUINEA-18 and MR766^MC^. Immunoblot assay on RIPA cell lysates were performed for the detection of the E protein from using anti-E mAb 4G2. *β*-tubulin served as a housekeeping protein.

**Figure S2.**
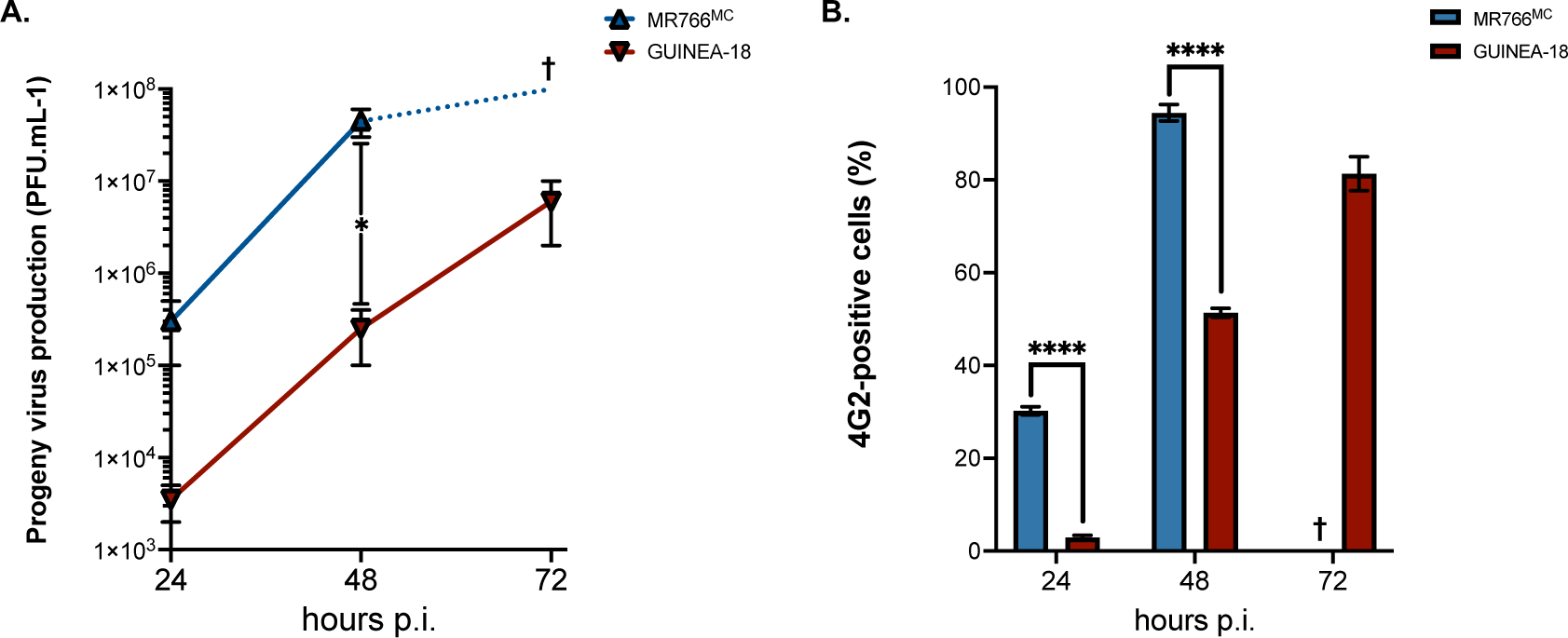
Infection of Vero cells with ZIKV at high multiplicity of infection. Vero cells were infected with MR766^MC^ and GUINEA-18 at the high m.o.i of 1. *In* (**A**), virus progeny production at various times post-infection. *In* (**B**), FACS analysis was performed on ZIKV-infected cells using anti-*pan* flavivirus E mAb 4G2 and the percentage of 4G2-positive cells was determined at various times post-infection. Asterisks indicate that the differences between experimental samples at each time point are statistically significant, using the unpaired *t* test (**** *P* < 0.0001, * *P* < 0.05).

**Figure S3.**
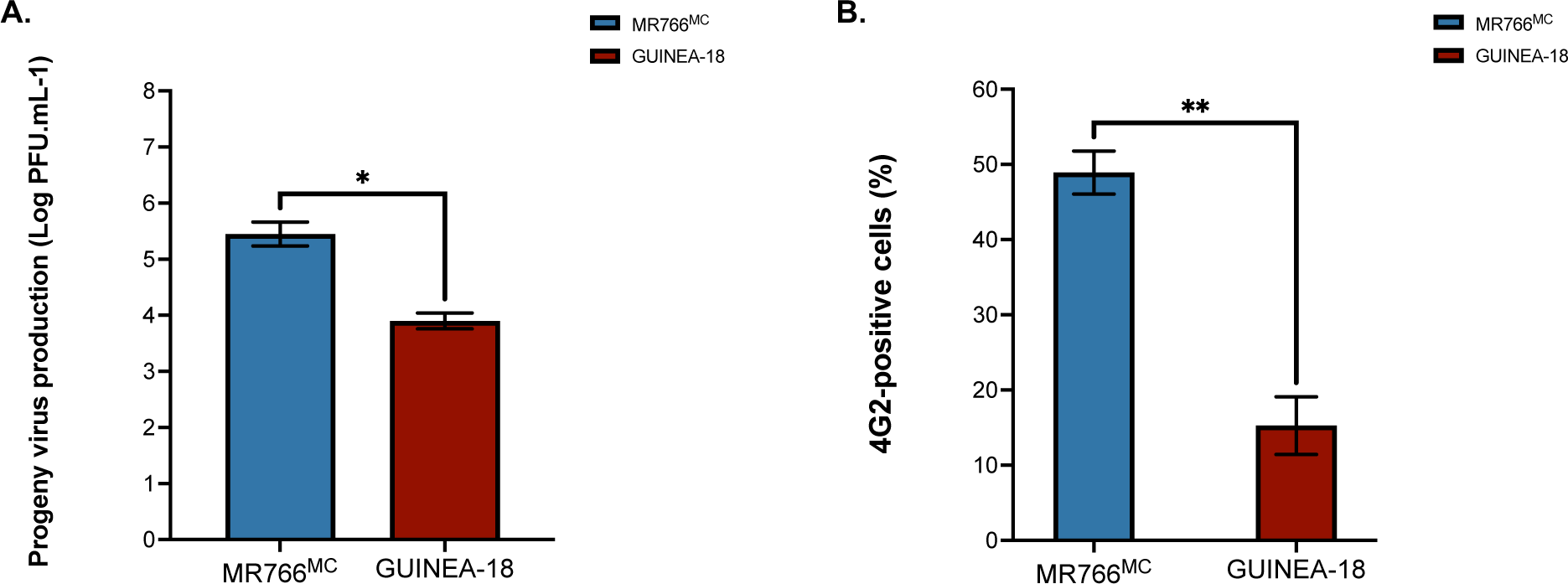
Infection of HCM3 cells with ZIKV. HCM3 cells were infected for 48 h with MR766^MC^ and GUINEA-18 at m.o.i. of 10. *In* (**A**), virus progeny production. *In* (**B**), FACS analysis was performed on ZIKV-infected cells using anti-*pan* flavivirus E mAb 4G2 and the percentage of 4G2-positive cells was determined. Asterisks indicate that the differences between experimental samples at each time point are statistically significant, using the unpaired *t* test (** *P* < 0.01; * *P* < 0.05).

**Figure S4.**
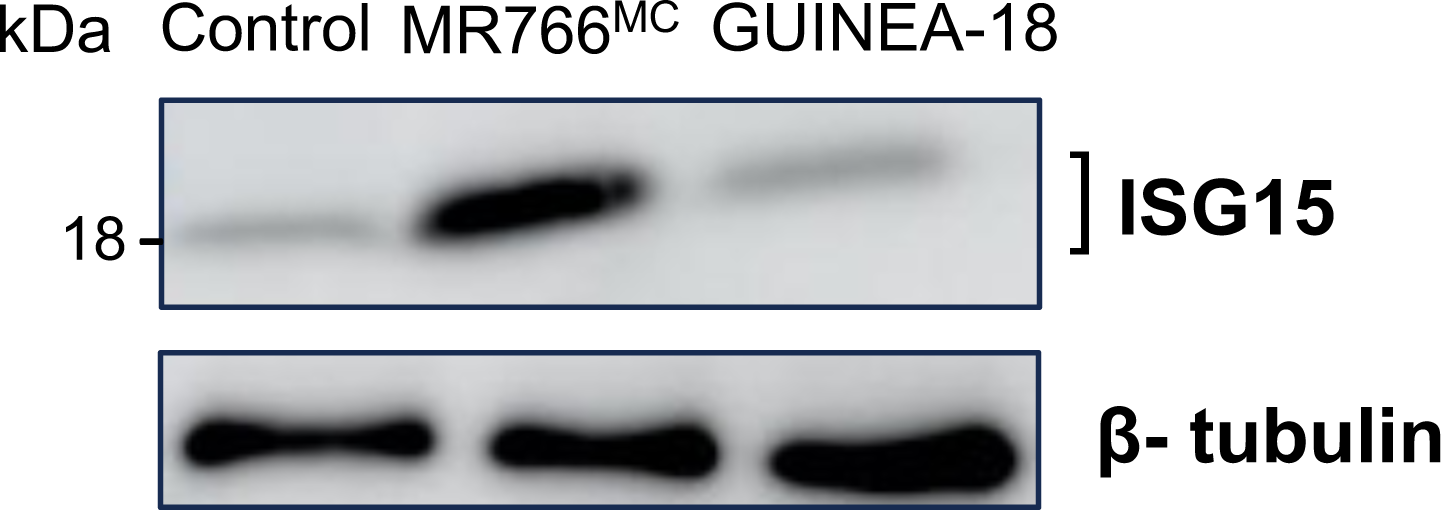
ISG15 expression in A549 cells infected by ZIKV. A549 cells were infected for 48h with MR766^MC^ or GUINEA-18 or mock-infected (control). Immunoblot assay on RIPA cell lysates was performed using anti-ISG15 mAb. *β*-tubulin served as loading protein.

**Figure S5.**
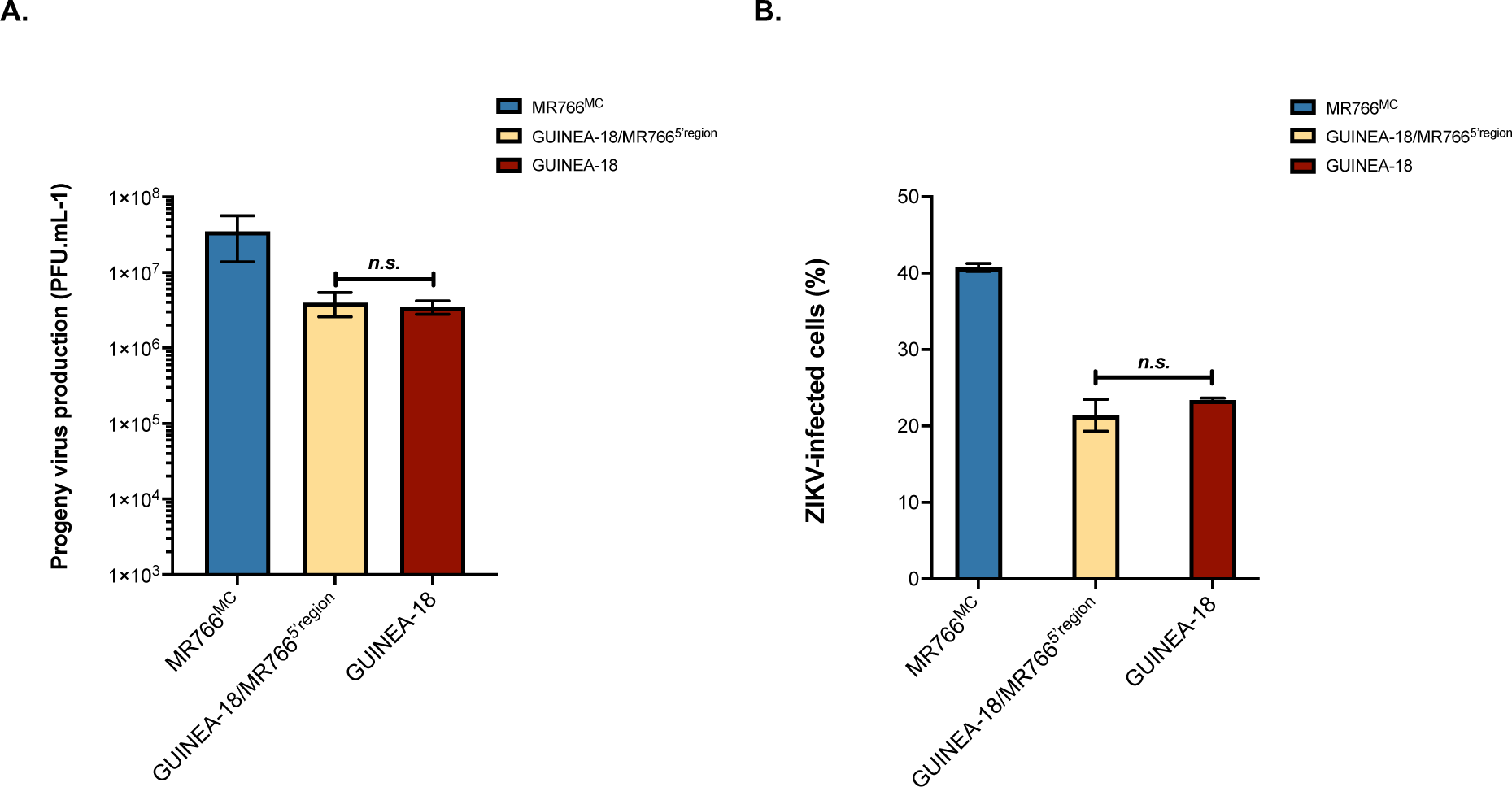
Replication of a GUINEA-18 chimera containing the first quarter of MR766 genomic RNA. A549 cells were infected for 48 h with GUINEA-18 or chimeric virus GUINEA-18/MR766^5’region^ at the m.o.i. of 1. Infection with MR766^MC^ served as a control. *In* (**A**), virus progeny production. *In* (**B**), FACS analysis was performed with anti-pan flavivirus E mAb 4G2. The percentage of 4G2-positive cells was determined. The results are the mean (± SEM) of two or three independent experiments. The differences between experimental samples at each time point are not statistically significant (n.s.), using the unpaired *t* test.

**Figure S6.**
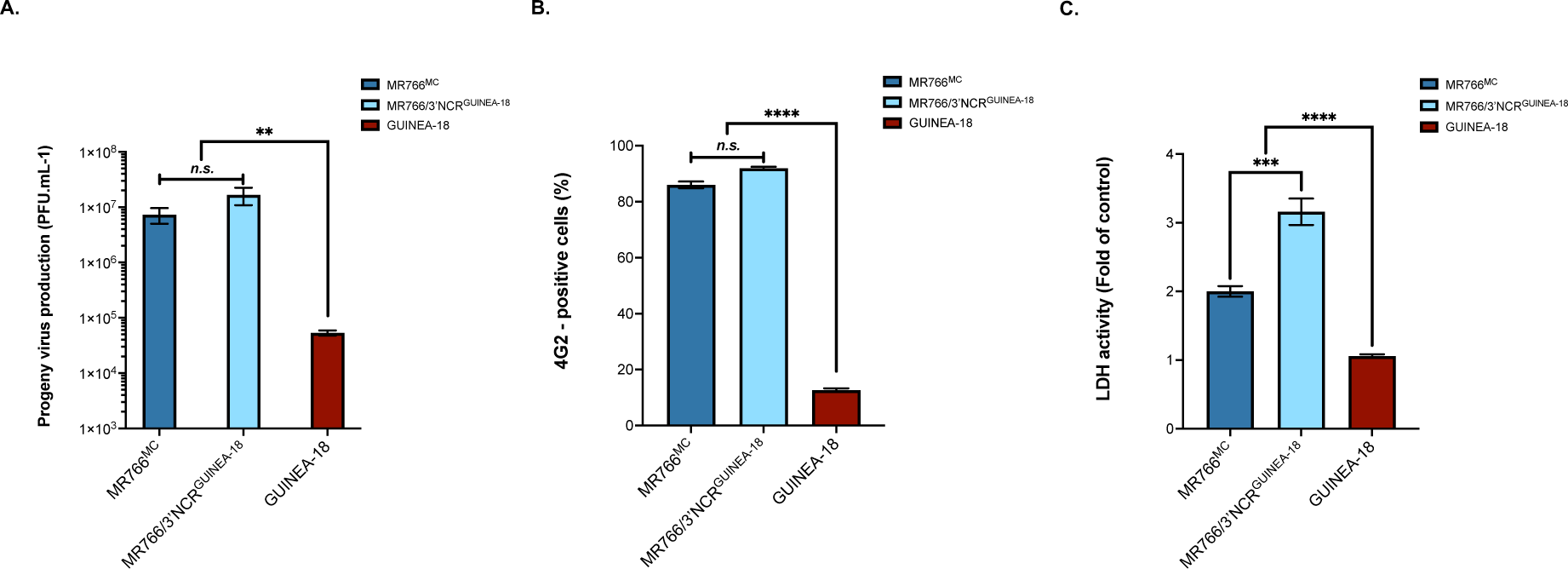
Replication of a MR766^MC^ chimera containing the 3’NCR from GUINEA-18. Vero cells were infected for 48 h with MR766/3’NCR^GUINEA-18^ chimera or parental viruses at m.o.i. of 0.1. *In* (A), virus progeny production. *In* (B), FACS analysis was performed with anti-E mAb 4G2. *In* (C), LDH activity was measured at 72h p.i. The results are the mean (± SEM) of two independent experiments. Asterisks indicate that the differences between experimental samples at each time point are statistically significant, using the unpaired *t* test or ANOVA *P* < 0.0001; *** *P* < 0.001; ** *P* < 0.01, *n.s*.: not significant).

**Figure S7.**
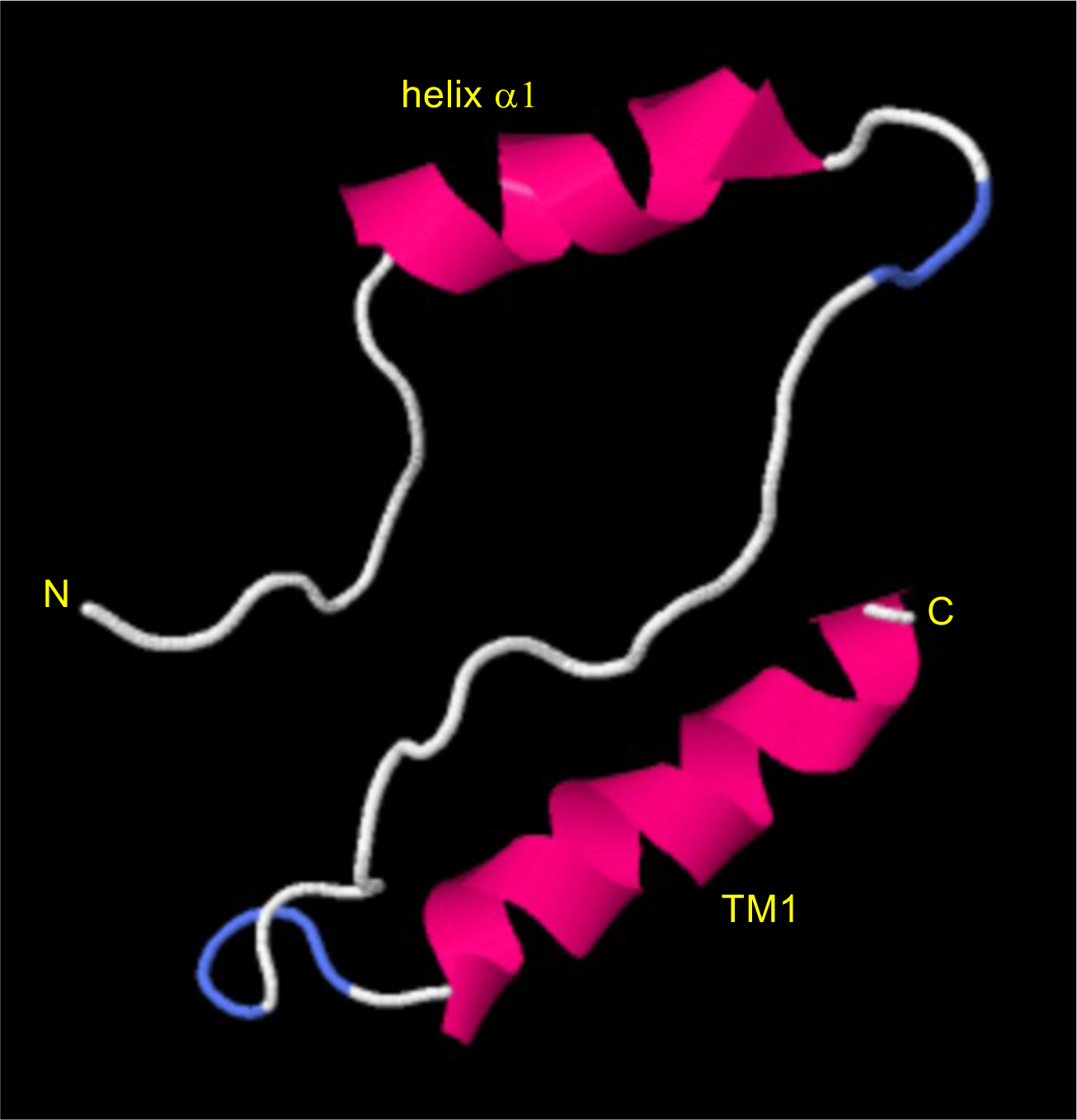
Three-dimensional prediction of the first 53 residues of ZIKV-15555 NS4B protein. The three-dimension structure prediction server Phyre^2^ was used to predict the 3D structures of the N-terminal region followed by transmembrane helix (TM1) of ZIKV-15555 NS4B protein. The 3D viewing of the predicted structure was performed using the JSmol molecular visualization system. The clusters of residues GWLETRTKSDIAHLM (NS4B-3/17) and PASAWAIYAALTTLI (NS4B-36/50) have propensity for forming a-helical structure (helix a1) and TM1, respectively. The central disordered structure corresponding to the cluster of GRKEEGTTIGFSMDIDLRP (NS4B-18/35) residues includes the three mutations at positions 20/24/26 that differentiate ZIKV-15555 from MR766.

**Figure S8.**
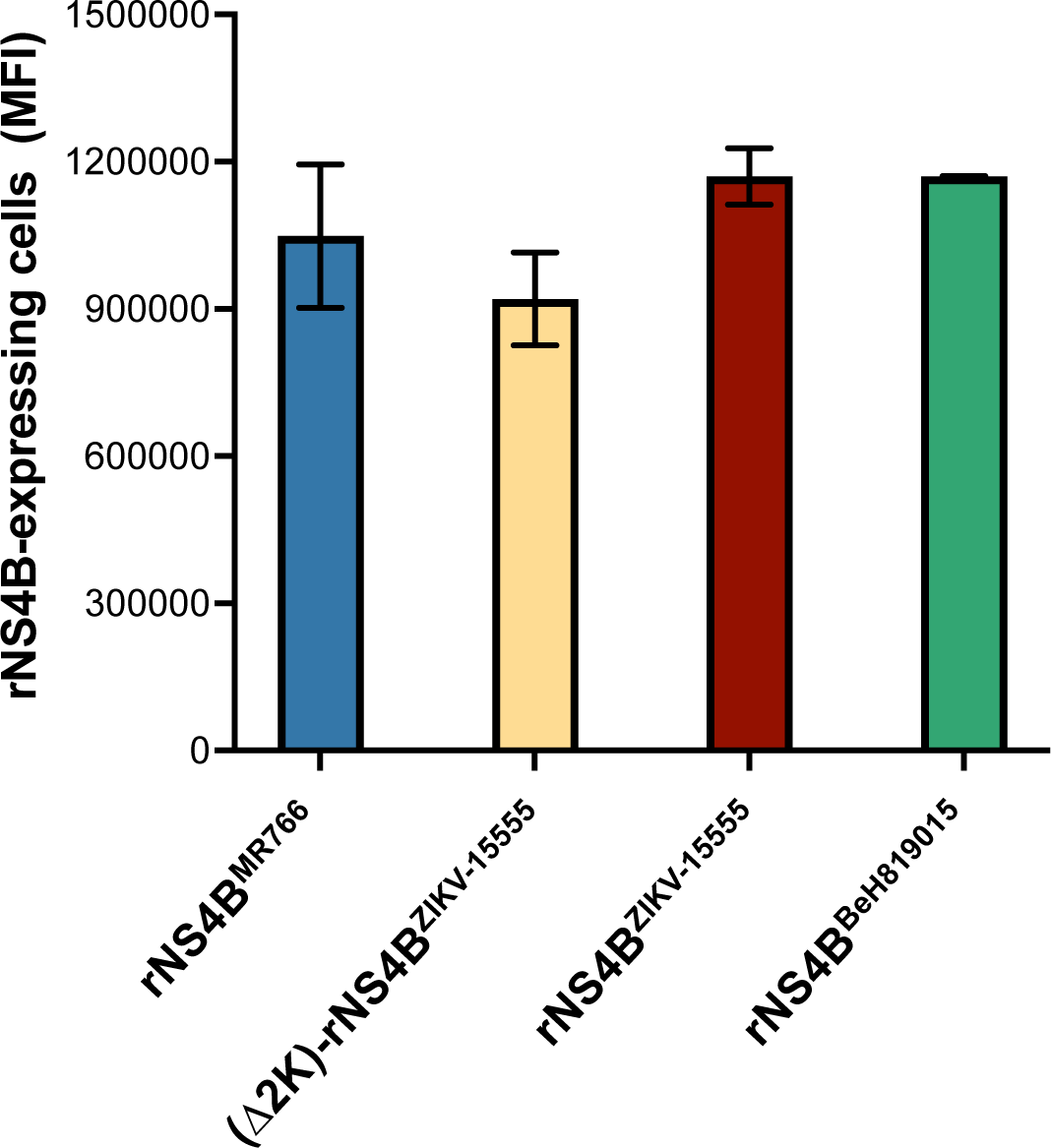
Expression level of rNS4B proteins. A549 cells were transfected for 24 h with plasmids expressing NS4B (rNS4B) protein from different ZIKV strains. FACS analysis was performed with anti-FLAG antibody and the mean of fluorescence intensity (MFI) of FLAG-positive cells was measured.

**Figure S9.**
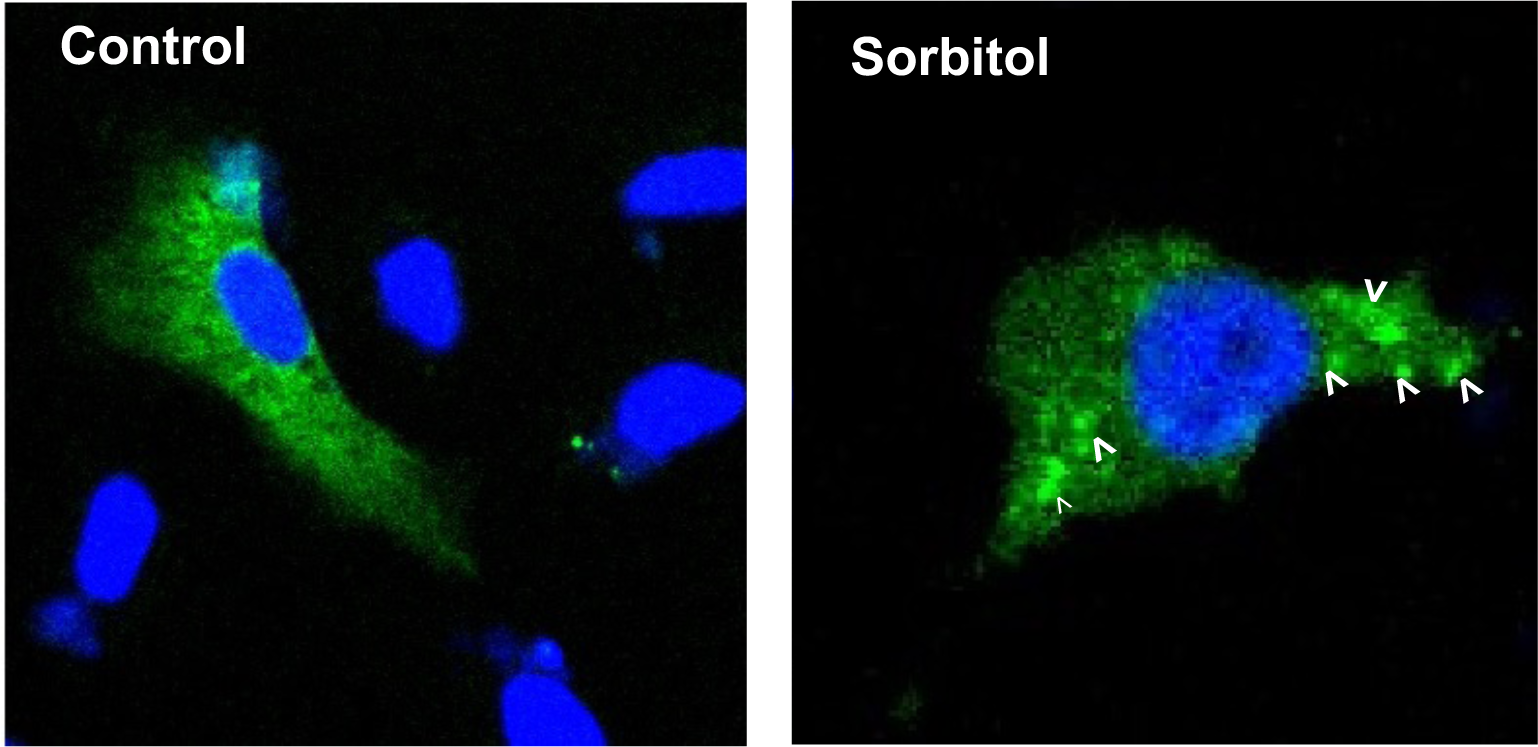
Stress granule formation in A549 cells subjected to a physiological stressor. A549 cells were transfected for 24 h with a plasmid expressing eGFP-G3BP fusion protein and then incubated with 0.4M sorbitol for 1.5 h (sorbitol) or mock-treated (control). Nuclei were stained with DAPI (blue). The cells were processed for confocal microscopy analysis. The blank arrow heads show the eGFP-positive condensates.

**Table S1.**
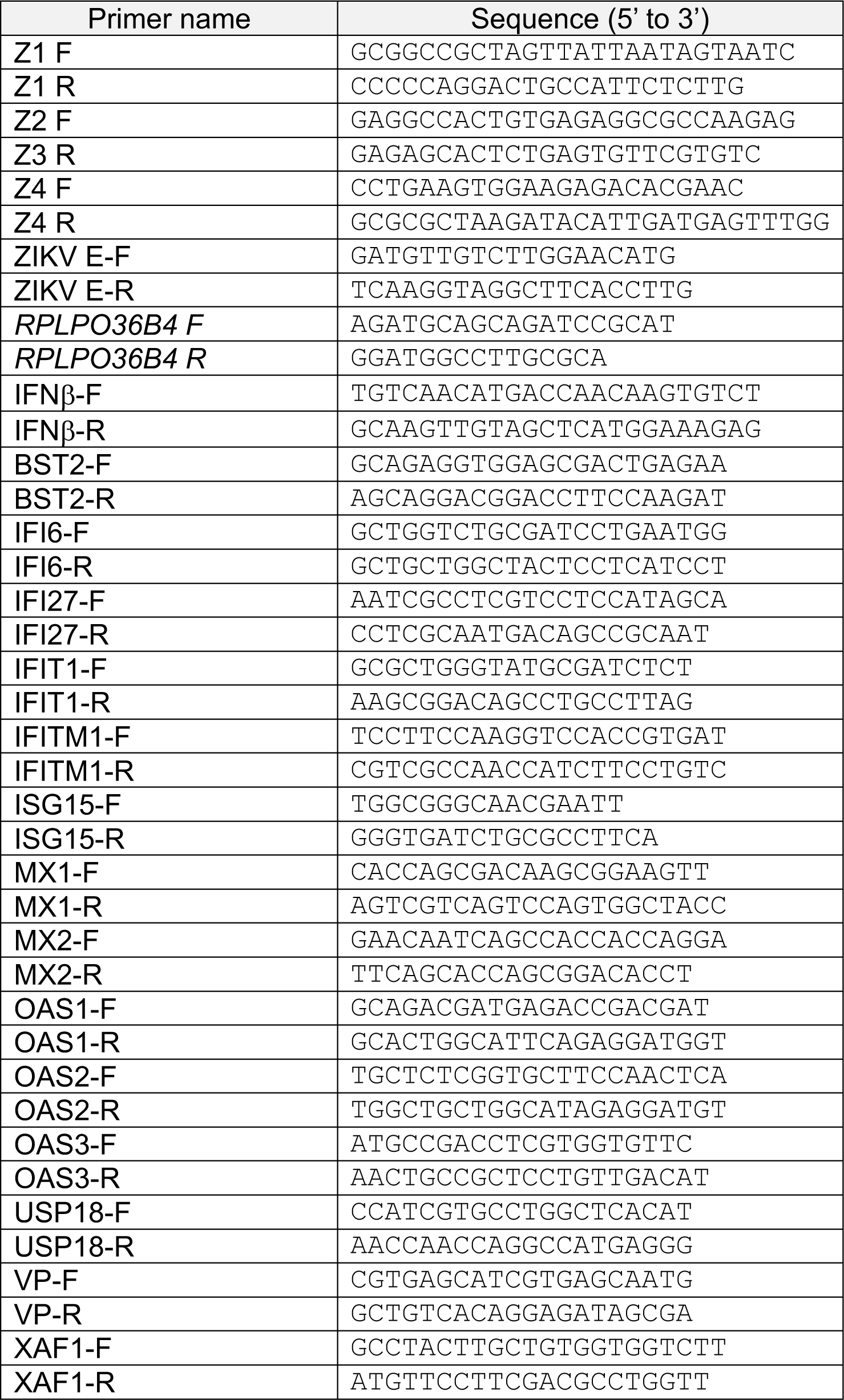
Sequences of primers for ISA method and RT-qPCR used in this study. Forward (F) and Reverse (R) primers.

Abbreviations.

Abbreviations
| Abbreviation | Value |
| --- | --- |
| Zika virus | ZIKV |
| Monkey kidney normal cells | Vero E6 |
| Human embryonic kidney cells | HEK-293T |
| Human carcinoma epithelial lung cells | A549 |
| Human microglial clone 3 cells | HCM3 |
| Ribonucleic acid | RNA |
| Non-coding regions | NCRs |
| Stem-loops | SLs |
| Nonstructural proteins | NS |
| Methyltransferase | MTase |
| Endoplasmic reticulum | ER |
| Viral replication complexes | VRCs |
| Infectious Subgenomic Amplicons | ISA |
| Dulbecco's modified Eagle's medium | DMEM |
| Fetal calf serum | FBS |
| Monoclonal antibody | mAb |
| Immunoglobulin G | IgG |
| Nitrocellulose | NC |
| Paraformaldehyde | PPFA |
| Bovine serum albumin | BSA |
| Green Fluorescent Protein | GFP |
| Enhanced Chemiluminescence | ECL |
| Laser scanning confocal microscope | LSM |
| Paraformaldehyde | PPFA |
| Lactate dehydrogenase | LDH |
| Glycan loop region | GL |
| Post-infection | p.i. |
| Multiplicity of infection | m.o.i. |
| Interferon | IFN |
| Interferon-Stimulated Genes | ISGs |
| 5' non-coding region | 5'NCR |
| Recombinant NS4B protein. | rNS4B |
| Enhanced Green Fluorescent Protein | eGFP |
| Stress granules | SGs |
| Interferon-stimulated genes | ISGs |
| RasGAP SH3 domain-binding protein | G3BP |

